# Longitudinal characterization of phenotypic profile of T cells in chronic hepatitis B identifies immune markers predicting HBsAg loss

**DOI:** 10.1101/2020.11.16.382838

**Authors:** Shue Xiong, Dan Zhu, Boyun Liang, Mingyue Li, Wen Pan, Junyi He, Hua Wang, Kathrin Sutter, Ulf Dittmer, Mengji Lu, Di Liu, Dongliang Yang, Jia Liu, Xin Zheng

## Abstract

The current desirable endpoint of treatment against chronic hepatitis B virus infection (cHBV) is to achieve a functional cure, which is defined as HBsAg loss (sAg-L) with or without anti-HBs seroconversion. However, the immunological features that are associated with functional cure have not been studied in detail. Here, a total of 172 cHBV patients including 31 sAg-L patients, and 24 healthy individuals were examined for their T cell phenotypic profile and HBV-specific T cell responses by flow cytometry. sAg-L patients showed distinct CD4 and CD8 T cell phenotype fingerprints compared to those of HBsAg-positive patients, as indicated by the upregulation of CD25, CD40L and CTLA-4 expression on CD4 T cells; HLA-DR, CD95 and PD-1 on both CD4 and CD8 T cells; as well as a potent HBcAg-specific CD8 T cell response. The changes in the T cell phenotype in sAg-L patients began during rapid HBsAg decrease upon treatment onset, were maintained after sAg-L. HLA-DR expression on T cells was positively correlated with the level of HBsAg reduction and the magnitude of the HBcAg-specific T cell responses in cHBV patients. Importantly, increased HLA-DR and CTLA-4 expression on CD4, as well as HLA-DR and TIM-3 expression on CD8 T cells were identified as predictive factors for HBsAg loss within 48 weeks of therapy in cHBV patients. The onset of HBsAg decrease and subsequent loss in cHBV patients on treatment is associated with significant alterations of both CD4 and CD8 T cell phenotypes. Characterization of the T cell phenotype in cHBV patients possessed greater predicative value for sAg-L.

## INTRODUCTION

The World Health Organization has implemented a Global Strategy on Viral Hepatitis to declare that the elimination of hepatitis B virus (HBV) is possible ^1^. Currently, most clinical practice guidelines recommend that the optimal endpoint of chronic HBV infection (cHBV) treatment is to achieve functional cure, which is defined as HBsAg seroclearance with or without anti-HBs seroconversion ^2-4^. This is based on the observation that the HBsAg seroclearance is associated with a beneficial effect on disease progression and reduced risk of hepatocellular carcinoma (HCC) development ^5,6^. Spontaneous HBsAg seroclearance is common in acute HBV infection in adults, but it is a rare clinical event in untreated chronic HBV infected patients with annual clearance rates ranging from 0.7% to 2.26% ^6,7^.

T cells are believed to play an irreplaceable role in HBV clearance and determine the outcome of HBV infection as demonstrated in the HBV chimpanzee model ^8,9^. Therefore, T cells from patients at different stages of HBV infection have been intensively characterized for their phenotypic profile and function ^10-13^. Available data indicate that the quantity and function of HBV-specific T cells are associated with the outcome of HBV infection. Strong HBV-specific CD4 and CD8 T cell responses were frequently detected in patients who spontaneously cleared the virus during acute HBV infection, but were not found in cHBV patients ^14,15^. During cHBV infection, an association between increased HBcAg/HBeAg-specific T cell responses and HBeAg seroconversion was also reported ^16^. Moreover, the recovery of HBV-specific T cell responses was observed in cHBV patients after long-term effective therapy with nucleos(t)ide analogues (NUCs), and in patients who achieved a functional cure either spontaneously or under antiviral treatment ^17^. A recent study also demonstrated that the presence of functional HBV-specific T cells may serve as a potential immune system biomarker to safely discontinue NUC therapy in CHB patients ^18^. Thus, a better understanding of HBV-specific T cell biology is believed to have significant implications for guiding clinical practice in cHBV treatment and for developing effective immunotherapy to cure cHBV. However, current available data on T cell responses related to cHBV functional cure mainly come from cross-sectional studies, and a longitudinal analysis of the T cell phenotype and function during the course of HBsAg loss (sAg-L) and seroconversion in cHBV patients is still lacking. In this study, T cell phenotype profiles and HBV-specific T cell responses were longitudinally analyzed in a group of cHBV patients prior to, during and after achieving functional cure upon treatment and compared with HBsAg-positive patients. Furthermore, T cell features correlating with rapid HBsAg decrease, loss and seroconversion in cHBV patients were characterized.

## RESULTS

### Alteration of the T cell phenotype in sAg-L patients and patients experiencing rapid HBsAg decrease

First, we characterized *ex vivo* whether immune cell populations and the T cell phenotype in PBMCs changed in sAg-L patients compared to HBsAg retained (sAg-R) patients and healthy controls by flow cytometry (as depicted in figure S1). No significant differences in absolute numbers and frequencies of B cells, monocytes, and dendric cells (DCs) were observed between sAg-L, sAg-R patients, and healthy controls (HC). The absolute numbers of total T cells as well as CD4^+^ T cells in sAg-R and sAg-L patients were significantly lower than those in HC. sAg-R but not sAg-L patients also showed significantly lower frequencies of total T cells as well as absolute numbers of CD8^+^ T cells than HC. sAg-L patients showed significantly higher CD8^+^ T cell frequencies than sAg-R patients and HC, and lower CD4^+^ T cell frequencies than HC (Figure S1C). Next, we analyzed the T cell phenotype by staining cell surface and intracellular markers associated with T cell activation (CD69, HLA-DR, CD95, CD25 and CD40L), exhaustion (PD-1, TIM-3 and CTLA-4) and effector function (granzyme B and CD107a). As shown in figure 1A, both CD4 and CD8 T cell phenotype fingerprints of sAg-L patients were distinct from those of sAg-R patients and HC. Statistical analysis of the percentage of cells expressing each marker revealed that CD4^+^ T cells of sAg-L patients had significantly increased expression of the activation markers HLA-DR, CD25 and CD95 compared to those from sAg-R patients and HC (Figure 1B). HLA-DR expression on CD8^+^ T cells in sAg-L patients was also significantly increased compared to sAg-R patients and HC (Figure 1C). Consistent with previous reports ^15,19,20^, the expression of the T cell exhaustion marker PD-1 on both CD4^+^ and CD8^+^ T cells in sAg-R patients was significantly higher than in HC. Interestingly, sAg-L patients showed a further increase in PD-1 expression on both CD4^+^ and CD8^+^ T cells compared to sAg-R patients. The CD40L and CTLA-4 expression on CD4^+^ T cells and CD95 expression on CD8^+^ T cells in sAg-L patients were significantly higher than in sAg-R patients, but not in HC (Figure 1B and 1C). No significant differences in CD69, Tim-3, granzyme B, and CD107a expression on both CD4^+^ and CD8^+^ T cells were observed between sAg-L, sAg-R patients, and HC (Figure S2). Moreover, no significant differences in CD4^+^ and CD8^+^ T cell phenotypes were observed between sAg-L patients with or without seroconversion to anti-HBs (Figure S3).

**Figure 1.**
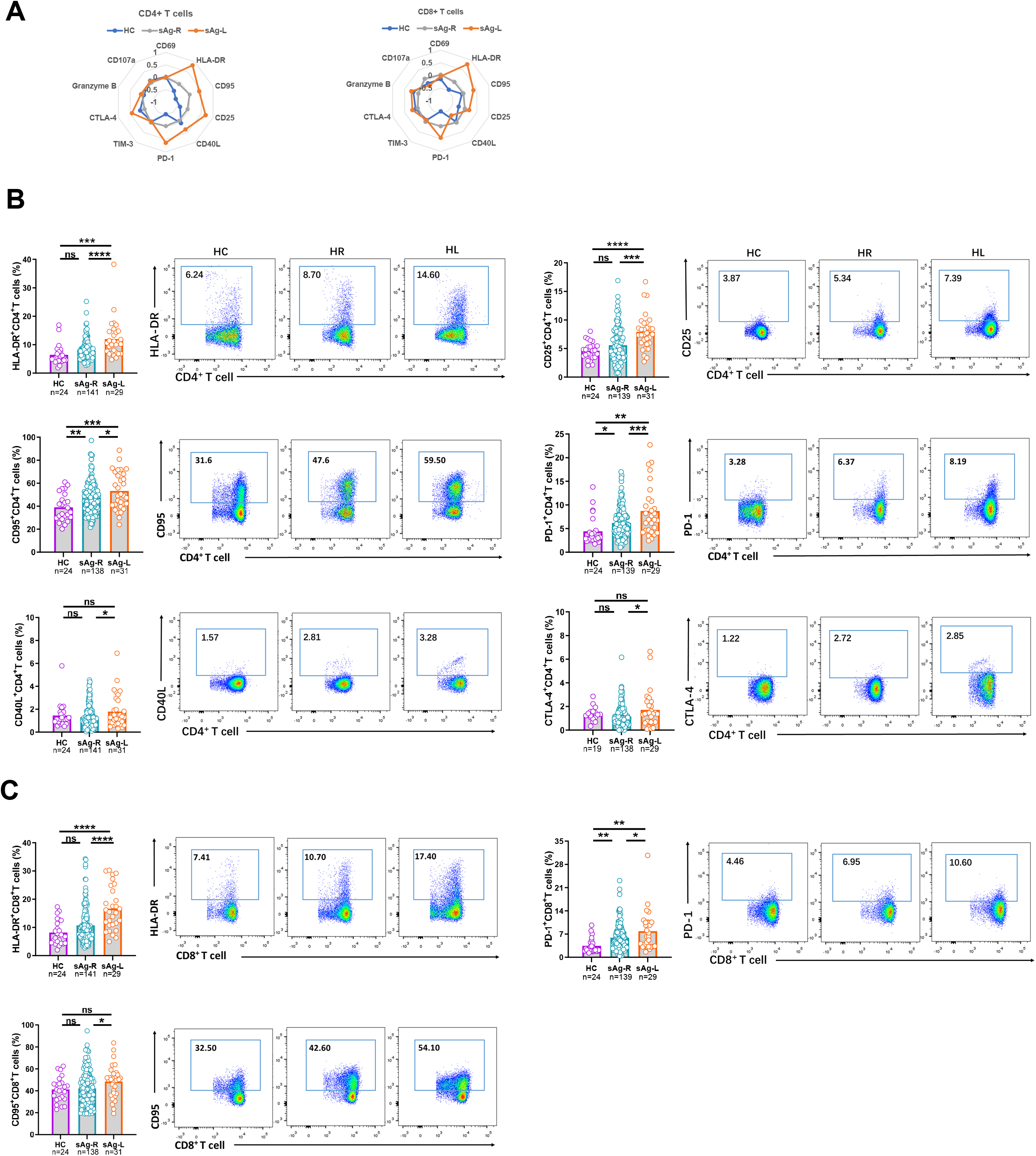
Characterization of T cell phenotype profiles in cHBV patients with HBsAg loss. (A) The mean percentage of CD4 and CD8 T cells expressing different T cell surface markers of healthy controls (HC), HBsAg retained patients (sAg-R) and HBsAg loss (sAg-L) were calculated by z-score and are depicted by radar plots. (B) Expression of HLA-DR, CD25, CD95, PD-1, CD40L, and CTLA-4 on CD4 T cells of HC, sAg-R and sAg-L patients were analyzed by flow cytometry. (C) Expression of HLA-DR, CD95, and PD-1 on CD8 T cells of HC, sAg-R and sAg-L patients were analyzed by flow cytometry. Error bars, mean ± s.e.m.; *P < 0.05; **P < 0.01; ***P < 0.001; ****P < 0.0001. Abbreviation: ns, not significant.

It has been shown that a lower HBsAg level is associated with a higher rate of HBsAg loss ^21,22^. Therefore, we subsequently examined whether cHBV patients experiencing a rapid HBsAg decrease (sAg-RD) also had an alteration of the T cell phenotype compared to cHBV patients with no decrease of serum HBsAg levels (sAg-ND) during the observational period. Nineteen patients in the sAg-R group experienced more than 30% decrease in HBsAg levels from the baseline within 6 months (sAg-RD30), while 30 sAg-R patients showed no decrease of HBsAg levels (Table S4). As shown in Figure 2, sAg-RD30 patients showed significant increases in HLA-DR, Tim-3 and CD107a expression on both CD4^+^ and CD8^+^ T cells compared to sAg-ND patients. CD40L and CTLA-4 expression on CD4^+^ T cells as well as CD69 expression on CD8^+^ T cells in sAg-RD30 patients were also significantly higher than those in sAg-ND patients (Figure 2A). No significant differences in CD25, CD95, PD-1, and granzyme B expression on T cells were observed between the two groups of patients (Figure S4). The differences in the T cell phenotype became even more evident when we examined a cohort of patients with a more profound HBsAg decrease (greater than 50% decrease, sAg-RD50). sAg-RD50 patients showed significant increases in HLA-DR, CD95, CD25, TIM-3, CTLA-4, and CD107a expression on CD4^+^, as well as HLA-DR and TIM-3 expression on CD8^+^ T cells compared to sAg-ND patients (Figure 2A). No significant differences in CD69, CD40L, PD-1 and granzyme B expression on T cells were observed between the two groups of patients (Figure S4). Consistent with these results, the levels of HLA-DR expression on CD4^+^ and CD8^+^ T cells positively correlated with the ratio of HBsAg reduction, but not with the quantities of HBsAg, in cHBV patients (Figure 2B and S5B). In addition, CD95, CD40L, CTLA-4 and CD107a expression levels on CD4 T cells, as well as TIM-3 and Granzyme B on CD8^+^ T cells, were positively correlated with the ratio of HBsAg reduction in cHBV patients (Figure 2B).

**Figure 2.**
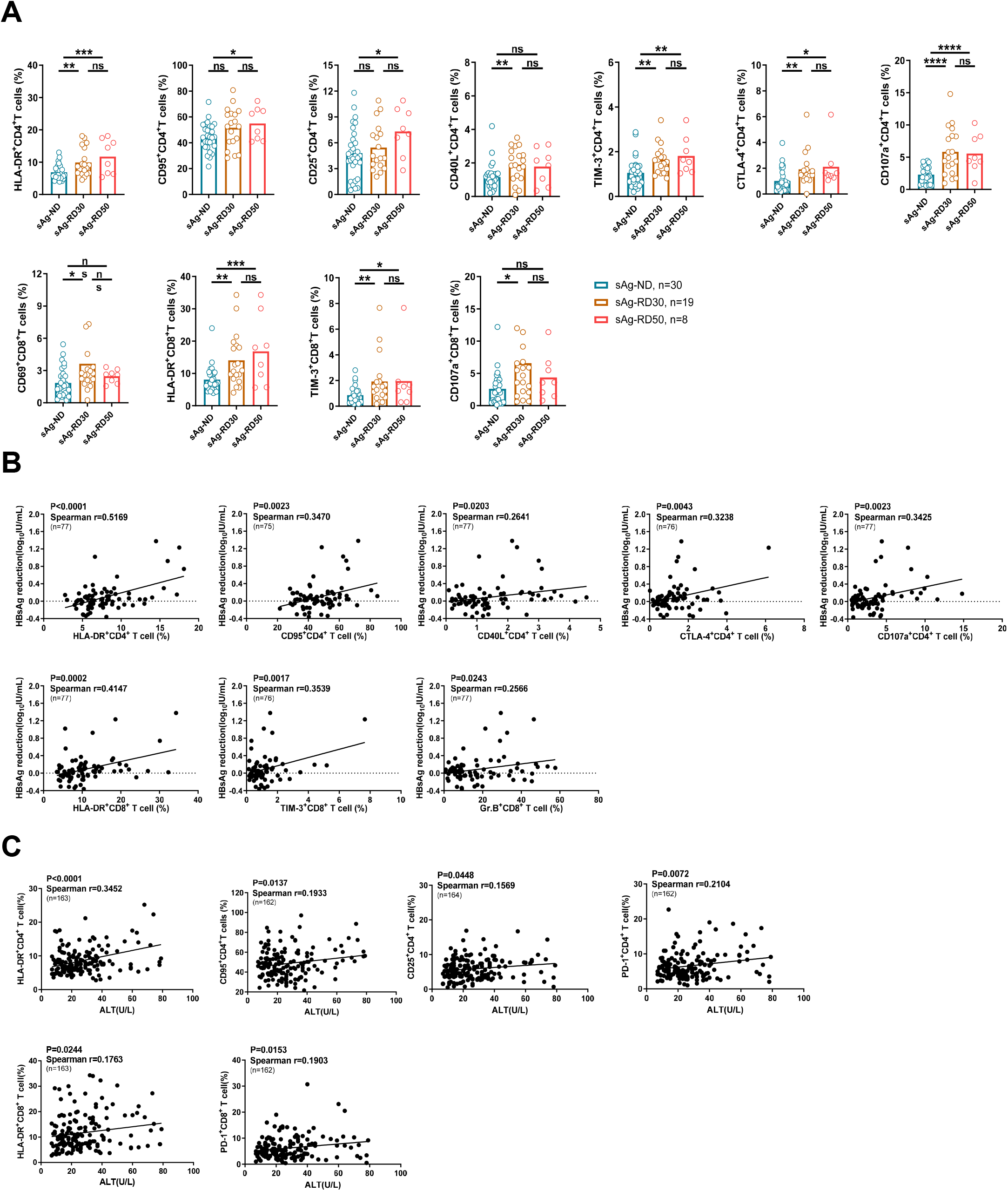
Characterization of T cell phenotype profiles in cHBV patients with HBsAg rapid decrease. (A) Expression of HLA-DR, CD25, CD40L, TIM-3, CTLA-4, and CD107a on CD4 T cells, and CD69, HLA-DR, TIM-3 and CD107a on CD8 T cells were analyzed by flow cytometry and were compared between cHBV patients who experienced more than 30% decrease of HBsAg levels (sAg-RD30, n=19), patients who experienced more than 50% decrease of HBsAg levels compared to the baseline within 6 months (sAg-RD30, n=8)and patients with no decrease of serum HBsAg levels (sAg-ND, n=30). (B) Pearson correlation analysis between expression of corresponding T cell markers and the extent of HBsAg reduction was performed in CHB patients. (C) Pearson correlation analysis between the expression of corresponding T cell markers and ALT levels was performed in CHB patients. Error bars, mean ± s.e.m.; *p<0.05, **p<0.01, ****p<0.0001, ns-not significant (p>0.05).

Next, we analyzed the correlation between the T cell phenotype and serum ALT and HBV-DNA levels. We found that CD95 and CD25 expression on CD4^+^ T cells, as well as HLA-DR and PD-1 expression on CD4^+^ and CD8^+^ T cells, were positively correlated with serum ALT levels (Figure 2C). No correlations between the T cell phenotype and serum HBV-DNA levels were observed (Figure S5D).

Overall, these data demonstrated that cHBV patients experiencing a rapid HBsAg decrease or loss have altered T cell phenotypes compared to HBsAg retaining patients mainly associated with cell activation.

### Association of HBV-specific T cell responses with alteration of T cell phenotype in sAg-L patients

During the natural history of cHBV, the quantity and function of HBV-specific T cells are shown to correlate with HBV control ^23^. Thus, we next examined how HBV-specific CD4^+^ and CD8^+^ T cell response change in our patient cohorts and how this correlated with T cell phenotypes. The PBMCs from cHBV patients were stimulated for 10 days with overlapping peptide pools spanning the HBcAg or HBsAg to induce HBV-specific CD4^+^ and CD8^+^ T cell expansion. Subsequently, the frequencies of IFN-γ, TNF-α or IL-2-producing HBV-specific CD4 or CD8 T cells were detected by FACS. sAg-L patients showed significantly increased frequencies of IFN-γ, TNF-α, or IL-2-producing HBcAg-specific CD8^+^ T cells, but not CD4^+^ T cells, compared to sAg-R patients (Figure 3B). Although sAg-L patients that seroconverted to anti-HBs showed higher frequencies of IFN-γ, TNF-α, or IL-2-producing HBcAg-specific CD8^+^ T cells than non-seroconverters, the differences were not statistically significant (Figure S6A). We also observed that sAg-RD50, but not sAg-RD30 patients showed significant increased frequencies of IFN-γ-producing HBcAg-specific CD4 T cells compared to sAg-ND patients (Figure 3C). No significant increases in the frequencies of IFN-γ, TNF-α, or IL-2-producing HBsAg-specific CD8^+^ T cells were observed in sAg-L patients compared to sAg-R patients (Figure S7A). Moreover, the correlation between HBV-specific T cell responses and T cell phenotype was analyzed. We demonstrated that HLA-DR, TIM-3, CTLA-4 and granzyme B expression on CD4^+^ T cells was positively correlated with HBcAg-specific CD4^+^ T cell responses, while HLA-DR, CD95 and CD25 expression on CD8^+^ T cells positively correlated with HBcAg-specific CD8^+^ T cell responses (Figure 3D). These results suggest that the activated phenotype of T cells during rapid HBsAg decrease and loss is associated with increased HBcAg-specific CD8^+^ T cell responses.

**Figure 3.**
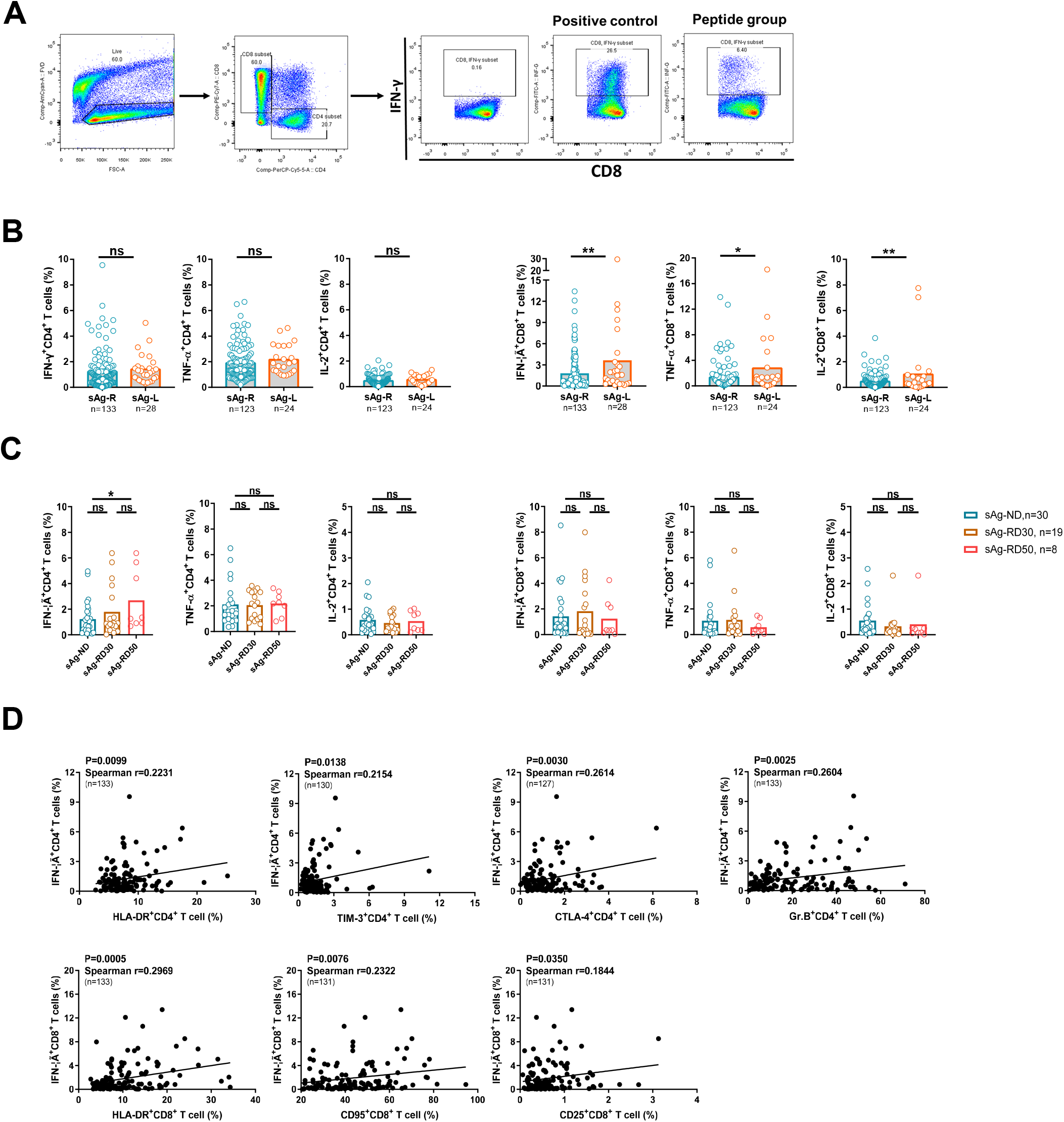
Characterization of HBV specific T cell responses in cHBV patients with HBsAg loss and rapid decrease. PBMCs from chronic hepatitis B patients were stimulated with overlapping peptide pools covering the entire sequences of HBcAg or HBsAg for 10 days. Cells were analyzed for IFN-γ, TNF-α, and IL-2 production by intracellular cytokine staining. (A) Gating strategy of intracellular cytokine staining analysis. (B) HBV-specific T cell responses were compared between HBsAg retained patients (sAg-R) and HBsAg loss (sAg-L) patients. (C) HBV-specific T cell responses were compared between sAg-ND, sAg-RD30 patients and sAg-RD50 patients. (D) Pearson correlation analysis between expression of corresponding T cell markers and frequencies of IFN-γ-producing HBcAg-specific T cells was performed in CHB patients. Error bars, mean ± s.e.m.; *p<0.05, **p<0.01, ****p<0.0001, ns-not significant (p>0.05).

### Alteration of the T cell phenotype in sAg-L patients is associated with IFN-α treatment

Accumulating clinical experience and increasing data from clinical trials have demonstrated that peg-IFN-α-containing therapy in cHBV significantly increased the rates of functional cure in selected cHBV patient cohorts ^24^. Our data also shows an increased percentage of patients receiving peg-IFN-α-containing therapy in the sAg-L group compared to the sAg-R group (58.1% vs 9.9%, Table S1). Therefore, we next examined whether the treatment strategy with IFN-α had an influence on the T cell phenotype. cHBV patients receiving peg-IFN-α-containing treatment (either alone or in combination with NUCs) showed significant increases in HLA-DR, CD95, CD25, CD40L, PD-1, TIM-3, and CTLA-4 expression on CD4 T cell, as well as HLA-DR, PD-1 and TIM-3 expression on CD8^+^ T cells, compared to patients receiving NUC treatment alone or no treatment (Figure 4A). Patients receiving peg-IFN-α-containing treatment also showed significantly higher frequencies of IFN-γ, TNF-α, or IL-2-producing HBcAg-specific CD8^+^ T cells than patients receiving NUC treatment alone (Figure 4B). These results suggest that alterations of the T cell phenotype in sAg-L patients is partially associated with peg-IFN-α treatment, which is known to exhibit immunomodulatory function and known to be beneficial for achieving HBsAg loss in cHBV patients.

**Figure 4.**
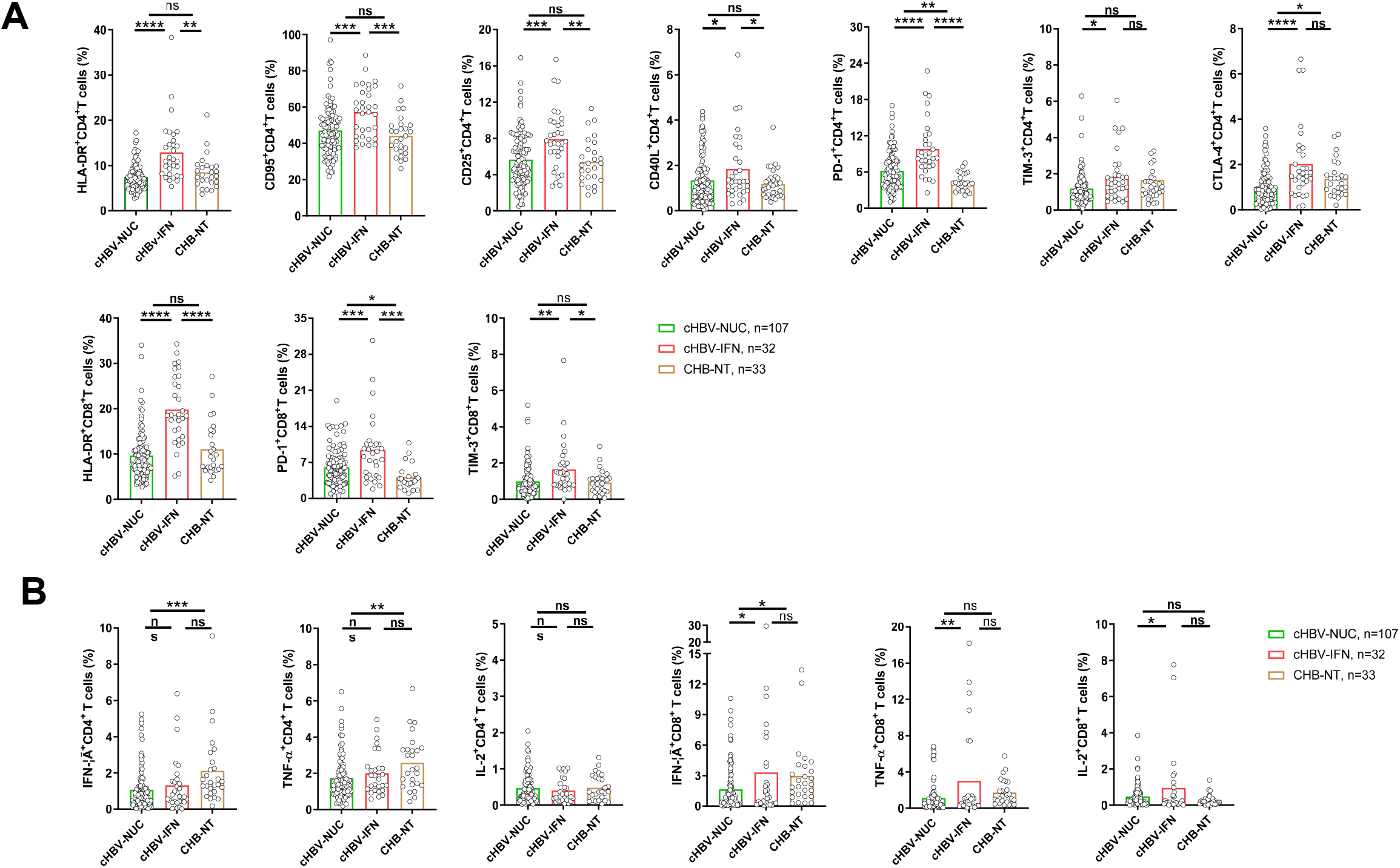
Characterization of T cell phenotype and HBcAg-specific T cell responses in cHBV patients received IFN-α treatment. T cell phenotypes (A) and HBcAg-specific T cell responses (B) were compared between cHBV patients received NUC monotherapy (cHBV-NUC), IFN-α contained therapy (cHBV-IFN), and no therapy (CHB-NT). Error bars, mean ± s.e.m.; *p<0.05, **p<0.01, ****p<0.0001, ns-not significant (p>0.05).

### Kinetics of the T cell phenotype during rapid HBsAg decrease, loss, and seroconversion

Next, we characterized the kinetics of the T cell phenotype alteration during rapid HBsAg rapid decrease and loss by analyzing the longitudinal data obtained from 7 sAg-ND, 5 sAg-RD50, and 6 sAg-L patients (Figure 5). These patients were followed for up to 60 weeks, monitored for virologic and clinical markers, and sampled 2-8 times for T cell phenotype and HBV-specific T cell response analysis. No significant differences in CD69, HLA-DR, CD25, CD40L, granzyme B, and CD107a expression on CD4^+^ and CD8^+^ T cells were observed between the sAg-ND, sAg-RD50, and sAg-L groups at baseline. Interestingly, the sAg-L group showed significantly higher CD95 and CTLA-4 expression on CD4^+^ T cells and TIM-3 expression on CD8^+^ T cells compared to the sAg-ND group at baseline (Figure 5A and 5B). The phenotypes of CD4 and CD8 T cells in the sAg-ND patient group remained relatively stable during the entire observational period. In contrast, the phenotypes of CD4^+^ and CD8^+^ T cells in both the sAg-RD50 and sAg-L patients group showed significant alterations during the follow-up period (Figure 5A and 5B). The expressions of HLA-DR, CD25, PD-1, and CTLA-4 on CD4^+^ T cells and HLA-DR on CD8^+^ T cells from sAg-RD50 patients was significantly higher than in sAg-ND patients during the follow-up, which was the period of rapid HBsAg reduction (Figure 5A and 5B). Consistently, significant increases in the expression of these markers were also observed in sAg-L patients both before and after HBsAg loss during the follow-up period. Furthermore, sAg-L patients showed significant upregulation of CD95 on CD4^+^ T cells and PD-1 on CD8^+^ T cells only before but not after HBsAg loss compared to sAg-ND patients (Figure 5A and 5B). An upregulation of CTLA-4 and TIM-3 on CD4^+^ T cells, as well as TIM-3 on CD8^+^ T cells was observed in sAg-L patients only before but not after HBsAg loss (Figure 5A and 5B). Next, the kinetics of the HBcAg-specific CD4^+^ and CD8^+^ T cell response were also analyzed. No significant differences in HBcAg-specific CD4^+^ and CD8^+^ T cell responses were observed between the three groups at baseline. The HBcAg-specific CD8^+^ T cell responses in sAg-ND and sAg-RD50 patients remained relatively stable during the entire observation period. sAg-L patients showed increases in HBcAg-specific CD8^+^ T cell responses only at later time points of the observation period (37-48 weeks of follow-up; Figure 5C). An increase of the HBcAg-specific CD8^+^ T cell responses in sAg-L patients was observed only before but not after HBsAg loss; however, the difference was not statistically significant (Figure 5C).

**Figure 5.**
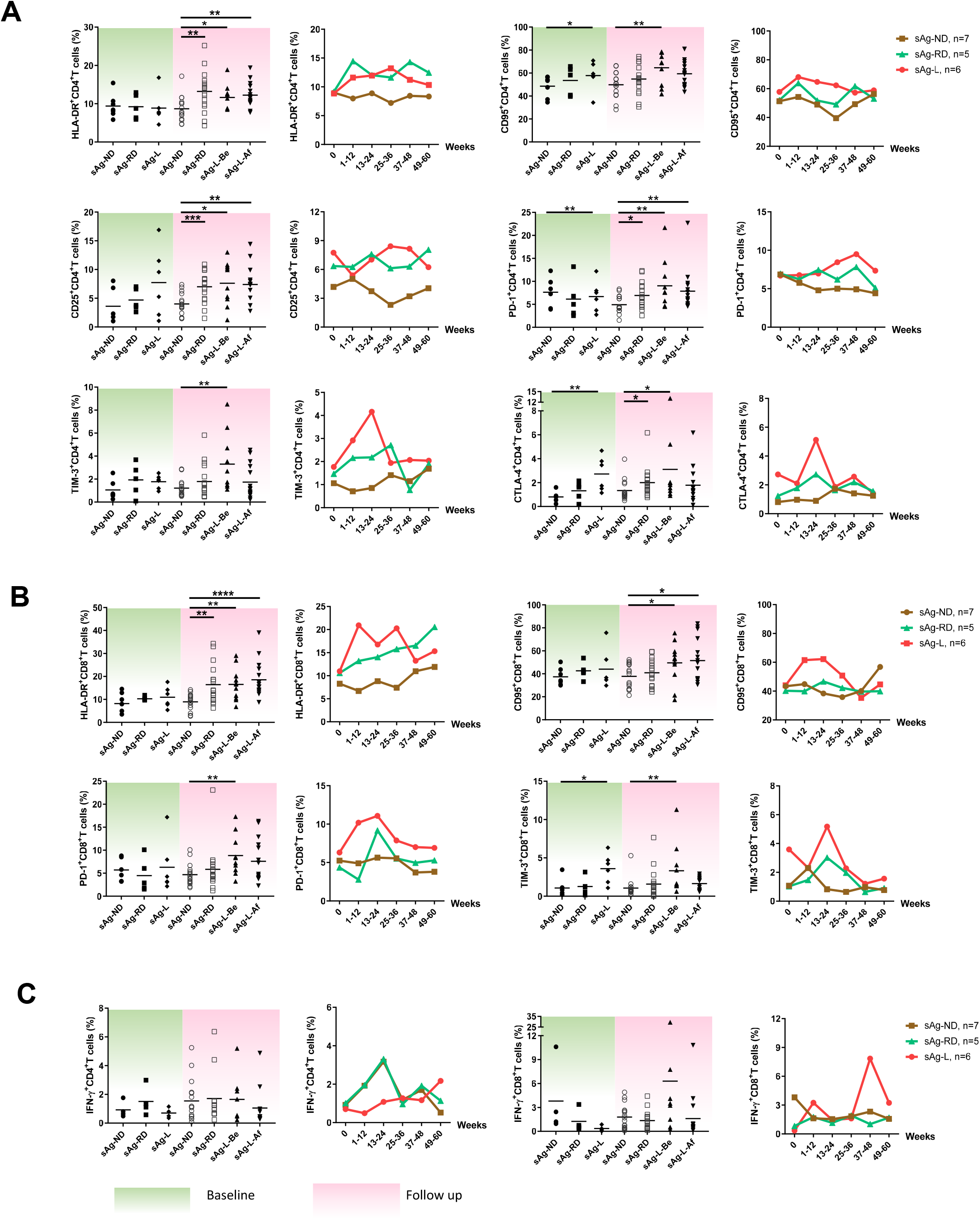
Kinetic analysis of T cell phenotypes and HBcAg-specific T cell responses in cHBV patients during HBsAg rapid decrease and loss. The phenotypes of CD4 T cells (A), CD8 T cells (B), and HBcAg-specific T cell responses (C) were longitudinally analyzed during the observation period. Left: Data were pooled and compared at either baseline (green background) and follow-up period (pink background) between cHBV patients with no HBsAg decrease (sAg-ND), HBsAg rapid decrease (sAg-RD50), and HBsAg loss (sAg-L). Right: Kinetic changes of T cell phenotype and HBcAg-specific T cell responses were demonstrated at indicated time points for sAg-ND, sAg-RD50, and sAg-L patients. Error bars, mean ± s.e.m.; *p<0.05, **p<0.01, ****p<0.0001, ns-not significant (p>0.05).

Next, the kinetics of T cell phenotype and serum HBsAg level were plotted individually for representative patients from each group. As shown in figure 6A, sAg-ND patients showed a minor alteration of the T cell phenotype and weak or undetectable HBcAg-specific CD8 T cell responses. An association of altered T cell phenotypes, as mainly represented by increasing HLA-DR expression on both CD4^+^ and CD8^+^ T cells, with a decrease in serum HBsAg levels was observed in sAg-RD50 patients. However, HBcAg-specific CD8^+^ T cell response was also barely detected in sAg-RD50 patients during rapid HBsAg decrease (Figure 6B). In sAg-L patients, an alteration of the T cell phenotype was observed during the course of HBsAg loss. Additionally, HBcAg-specific CD8^+^ T cell responses were more frequently detected in sAg-L patients during and after HBsAg loss (Figure 6C). The correlation of the T cell phenotype change with the HBsAb titer was also kinetically analyzed in 6 cHBV patients with seroconversion to anti-HBs. In most of the cases, the upregulation of HLA-DR expression on CD4^+^ and CD8^+^ T cells was accompanied by an increase in HBsAb levels (Figure S9D).

**Figure 6.**
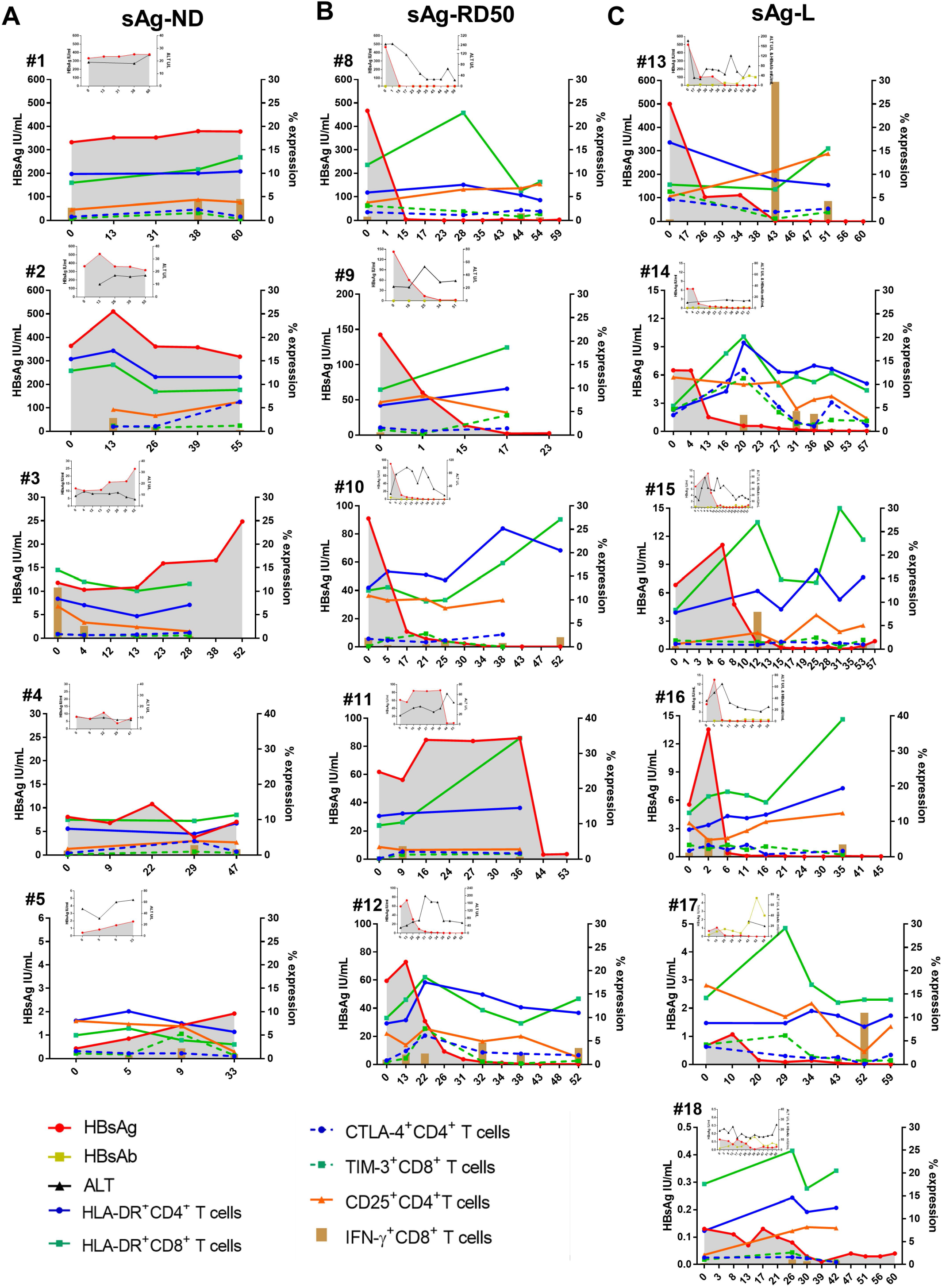
Kinetic analysis of T cell phenotypes and HBcAg-specific T cell responses in individual cHBV patients. (A) Data from 5 representative patients with no HBsAg reduction. (B) Data from 5 patients who experienced greater than 50% decrease of HBsAg levels than the baseline within 6 months. (C) Data from 6 patients who experienced HBsAg loss.

### Predicting HBsAg loss in cHBV patients by T cell surface markers

Our data demonstrate that the alteration of T cell phenotype might occur prior to HBsAg loss, therefore, we next evaluated the predictive performance of HLA-DR, CD25, CD95, PD-1, TIM-3, and CTLA-4 expression levels on T cells, as well as the frequencies of IFN-γ-producing HBcAg-specific CD8^+^ T cells in predicting HBsAg loss within 48 weeks of follow-up. This time frame was chosen, because all incidents of HBsAg loss in our patient cohort occurred within this period. The ROC curve analysis was performed in a total of 54 cHBV patients with HBsAg less than 600 IU/ml. The results showed that HLA-DR or CTLA-4 expression on CD4^+^ T cells, as well as HLA-DR, or TIM-3 expression on CD8^+^ T cells were significant predictors for 48-week HBsAg loss (Figure 7). The AUC of CD4^+^ T cell HLA-DR expression was 0.773 (p=0.031, [95% confidence interval, 0.607-0.939]), the optimal cutoff value was 7.35% (sensitivity=100% and specificity=55.30%). The AUC of CD4^+^ T cell CTLA-4 expression was 0.785 (p=0.024, [95% confidence interval, 0.608-0.963]), the optimal cutoff value was 1.49% (sensitivity=83.30% and specificity=70.20%) (Figure 7A). The AUC of CD8^+^ T cell HLA-DR expression was 0.807 (p=0.005, [95% confidence interval, 0.637–0.977]), the optimal cutoff value was 12.10% (sensitivity=83.33% and specificity=72.34%). The AUC of CD8^+^ T cell TIM-3 expression was 0.883 (p=0.002, [95% confidence interval, 0.705–1.000]), the optimal cutoff value was 3.26% (sensitivity=66.70% and specificity=97.90%) (Figure 7B). A previous study has reported that serum HBsAg level is a predictor for HBsAg loss ^22^. However, our result showed that the AUC of HBsAg for predicting 48-week HBsAg loss was only 0.135 (p=0.018, [95% confidence interval, 0.000-0.467]) (Figure 7C). Therefore, these results suggest that CD4^+^ T cell HLA-DR and CTLA-4 expression, and CD8^+^ T cell HLA-DR and TIM-3 expression possess a predictive value for 48-week HBsAg loss in cHBV patients.

**Figure 7.**
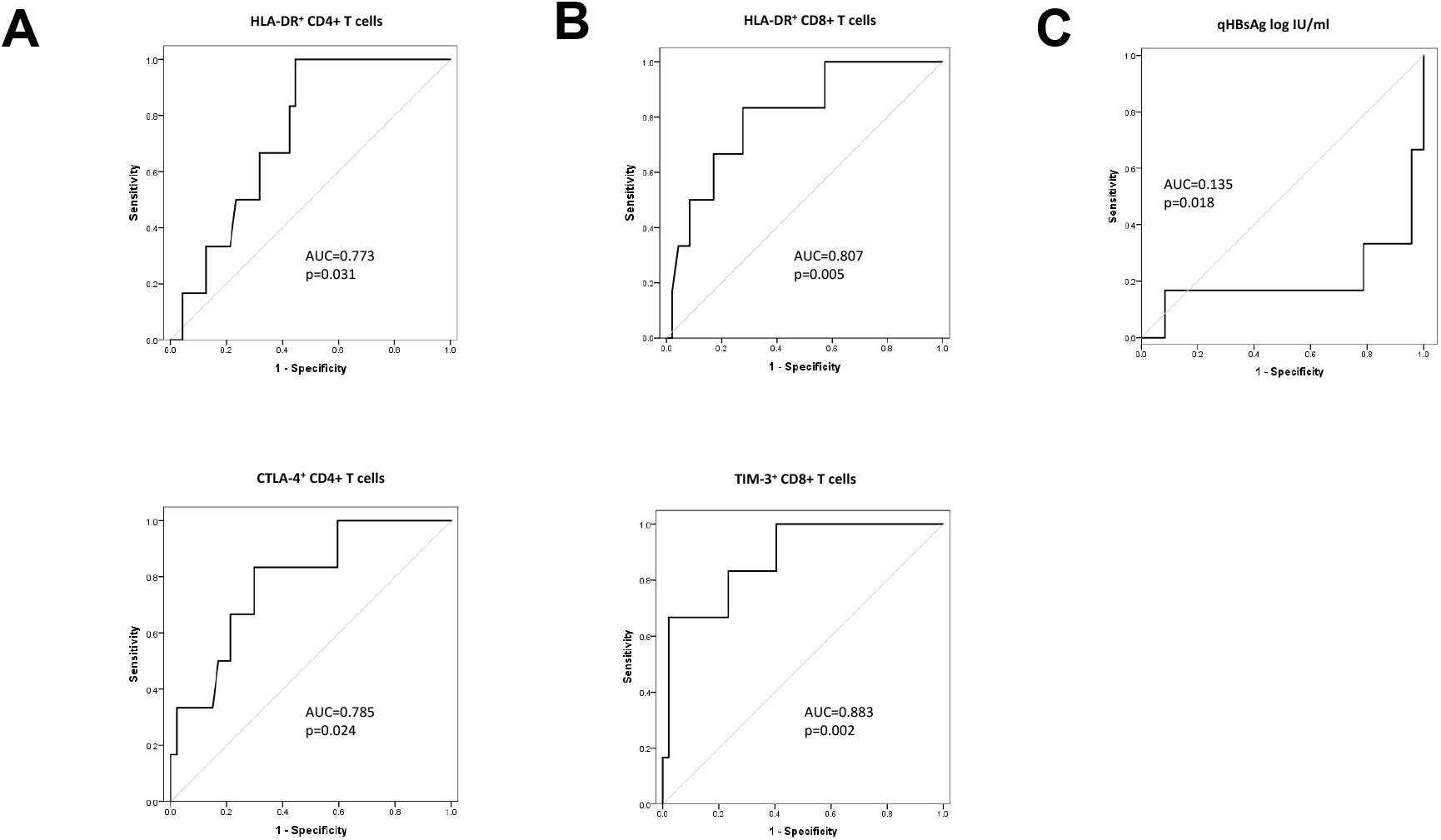
Predictive factors for HBsAg loss within 48 weeks in cHBV patients. ROC curve and AUC were calculated for HLA-DR and CTLA-4 expression of CD4 T cells (A), HLA-DR and TIM-3 expression of CD8 T cells (B) and HBsAg (C) by using R package “pROC”.

## DISCUSSION

In the current study, we longitudinally characterized T cell phenotype profile and HBV-specific response during HBsAg rapid decrease, loss and seroconversion. Importantly, all analyses were performed directly *ex vivo*, since freezing and thawing T cells before analysis may induce profound changes in their functionality ^18^. Our findings reveal that the courses of HBsAg rapid decrease and loss in cHBV patients is associated with significant alteration of both CD4 and CD8 T cell phenotypes. We identify that T cells upregulate the expression of a panel of surface molecules, including CD25, CD40L, and CTLA-4 expression on CD4 T cells, and HLA-DR, CD95, and PD-1 expression on both CD4 and CD8 T cells. The expression of HLA-DR, CD95, CD40L, CTLA-4, CD107a on CD4 T cells, and HLA-DR, TIM-3, granzyme B on CD8 T cells is positively corelated with the extent of HBsAg decrease. The expression of HLA-DR, TIM-3, CTLA-4, granzyme B on CD4 T cells and HLA-DR, CD95, CD25 on CD8 T cells were positively corelated with the intensity of HBcAg-specific T cell response. HLA-DR and CTLA-4 expression on CD4, as well as HLA-DR and TIM-3 expression on CD8 T cells were identified as predictive factors for HBsAg loss within 48 weeks in cHBV patients.

Functional cure is a rare clinical event in cHBV patients, with an annual clearance rate of 0.33% taking on average more than 50 years to clear the virus ^25,26^. Previous studies have shown that improved peg-IFN-α-containing treatment strategies, such as “adding-on” or “switching to” peg-IFN-α in patients who achieved long-term effective virological remission by NUCs, may significantly increase the HBsAg loss rates to more than 20%^24,27,28^. In line with these reports, an increased rate of receiving peg-IFN-α-containing therapy in sAg-L patients than sAg-R patients was observed in the patient cohort of this study. Peg-IFN-α has been shown to have both direct antiviral and immunomodulatory effects for the treatment of CHB, and it is conceivable that treatment outcome may be mostly triggered by the immunomodulatory effects of Peg-IFN-α on the innate and adaptive immune responses ^29^. The mechanisms responsible for the peg-IFN-α mediated restoration of anti-HBV immune functions are not fully understood. Treatment with peg-IFN-α in CHB patients has been shown to lead to a significant functional augmentation of NK cells, however, quite different effects were observed on T cells ^30^. IFN-α therapy led to a striking reduction of CD8+ T cells and showed no effect on the restoration of frequency and early effector functions of HBV-specific CD8+ T cells ^30,31^. In contrast to these findings, we observed significant increases in HBV-specific CD8 T cell responses in patients who received IFN-α treatment compared to those without. In addition, patients who received IFN-α treatment demonstrated a more active phenotype of global T cells than IFN-α untreated patients. This is probably due to the majority (> 70%) of the IFN-α treated patient in our study also received NUCs treatment, while all patients in previous studies ^30,31^ only received IFN-α monotherapy. It has been shown that HBV-specific CD8+ T-cell functions could be restored, at least in part, after long-term treatment with NUCs ^17^. Therefore, it is assumed that peg-IFN-α could improve the immunomodulatory action of NUCs and vice versa ^32,33^. Our observation supports this assumption and indicates that IFN-α therapy may also be favorable for the T cell arm of immune system to control HBV infection in long-term NUC treated patients.

Our data demonstrates that the courses of HBsAg rapid decrease and loss are associated with increasing global T cell activation as indicated by the upregulation of HLA-DR, CD25, and CD95 on T cells. This is in line with a recent report showing that global T cells of patients with subsequent HBsAg loss showed a more activated phenotype compared to patients with retained HBsAg ^34^. Interestingly, we observed that markers usually used to represent CD8 T cell exhaustion statues during chronic viral infection, such as PD-1 and TIM-3, were also upregulated on CD8 T cells during the course. Previous studies have shown that PD-1 is not a definitive marker for complete functional exhaustion but can also be expressed on at least partially functional T cells in the context of a chronic infection ^18,35-37^. Patients with partial immune control of HBV infection display higher levels of intrahepatic PD-1^+^ CD39^+^ tissue-resident CD8+ T cells that possess the capacity to mount immediate and strong cytokine responses ^38^. PD-1 has also been shown to prevent virus-specific CD8+ T cells from terminal exhaustion and to contribute to the survival of memory T cell populations ^39^. Interestingly, a recent study has demonstrated that almost all HBcAg- and HBV polymerase-specific CD8 T cells in cHBV patients were PD-1 positive ^40^. Similar results were observed by us in an HBV replication mouse model (data not shown). Taken together, the increases in frequencies of PD-1+ T cells during the course of HBsAg loss may indicate the expansion and activation of HBV-specific T cell populations, but not enhanced T cell exhaustion. Also, the upregulation of PD-1 and TIM-3 expression on T cells during acute-resolving HBV infection has been reported ^41-43^, and higher TIM-3 expression on CD8 T cells was found to be associated with elevated serum anti-HBs production in HBV-carrying mice ^44^. Together with the observation of increases in HBcAg-specific CD8 T cell response, our data suggest that T cells may experience similar course of activation during HBs loss and seroconversion in chronic HBV infection as do in acute HBV infection.

The presence of functional HBV-specific T cells is considered as a potential immunological biomarker to evaluate the immune status of cHBV patients and to guide clinical practice, such as safe NUC therapy discontinuation ^18^. However, the process of detecting HBV-specific CD8 T cell responses in cHBV patients is relatively complicated to be broadly applied for diagnostic purposes. In this study, we have identified that HLA-DR expression on CD8 T is positively correlated with the intensity of HBcAg-specific CD8 T cell response, and thus it might serve as an easy surrogate detection marker for HBcAg-specific CD8 T cell response in cHBV patients. Besides, we have also identified HLA-DR expression on both CD4 and CD8 T cells, as well as CTLA-4 expression on CD4 T cells and TIM-3 expression on CD8 T cells, as candidate immunological biomarkers for predicting HBsAg loss within 48 weeks. Previous studies have indicated that a lower HBsAg level and HBV DNA level at baseline and older age may predict HBsAg loss and seroconversion ^45,46^. Therefore, it would be interesting to investigate whether combining both virological and immunological biomarkers possess a greater power to predict HBsAg loss and seroconversion.

There are several limitations of the current study. First is the lack of access to hepatic tissue samples, and thus we cannot compare the phenotypes of intrahepatic T cells with their peripheral counterparts. However, previous reports have shown that the immunologic changes of T cells and NK cells in peripheral blood could closely mirror those in the liver in cHBV patients ^47,48^. Second is the lack of analyzing phenotypes of HBV-specific T cells. Due to the low frequency of HBV-specific T cells present in cHBV patients, phenotype analysis of such rare cell populations requires large volume of peripheral blood and applying a pMHC tetramer-based technic to enrich the cells ^35,40^. We were unable to perform such an analysis due to the limitation of sample volume we were allowed to take from patients. It would be important to compare whether the phenotype change of global T cells represents that of the HBV-specific T cells.

In the current study, we present the longitudinal characteristics of T cell phenotypes and responses during the course of HBsAg rapid decrease, loss and seroconversion in the peripheral blood of cHBV patients and identified potential immunological biomarkers to predict HBsAg loss. This knowledge may assist in developing assays to precisely evaluate immune status and to improve treatment strategies for achieving functional cure in cHBV patients.

## METHODS

### Subjects

A total of 24 healthy controls (HC), 141 HBsAg retained (sAg-R) cHBV patients and cHBV patients with 31 HBsAg loss (sAg-L) were recruited at the Department of Infectious Diseases, Union Hospital, Tongji Medical College, Huazhong University of Science and Technology from June 2016 to March 2019. The diagnosis and phase classification of chronic HBV infection were based on the EASL 2017 Clinical Practice Guidelines on the management of hepatitis B virus infection ^3^. All patients tested negative for HIV, HCV, hepatitis E virus, and hepatitis delta virus. Patients with alcoholic liver disease, autoimmune disease, malignancy, or serious illness of other systems were excluded. Informed consent was obtained from each patient, and the study protocol was approved by the local medical ethics committee of Union Hospital, Tongji Medical College, Huazhong University of Science and Technology in accordance with the guidelines of the Declaration of Helsinki. Twenty-four healthy volunteers were enrolled as controls.

### Characteristics of study cohort

Demographic profiles and detailed patient characteristics are listed in Table S1 and Table S2. All patients in the sAg-R group remained serum HBsAg positive during the observation period, and 12 of them had been longitudinally monitored for their T cell phenotype profile and HBV-specific T cell response for 60 weeks. In the sAg-L patient group, 6 patients were serum HBsAg positive at the start of the study and experienced HBsAg loss during the observation period, among 3 of them showed HBsAg seroconversion. Twenty-five patients were already serum HBsAg negative when enrolled and 13 of them showed HBsAg seroconversion during the observation period. There were no significant differences in age, ALT levels, and sex between the sAg-R and sAg-L patient groups. A significantly higher percentage of patients in the sAg-L group compared to the sAg-R group (58.07% vs 9.93%) had received interferon α (IFNα) treatment either alone or in combination with NUCs for more than 24 weeks. For cHBV patients, 107 of them received NUCs monotherapy treatment, 32 received interferon α (IFN-a) combined or monotherapy treatment and the others received no antiviral treatment.

### Peripheral blood mononuclear cells (PBMCs)

PBMCs of healthy controls and patients were isolated using Ficoll density gradient centrifugation (DAKEWE Biotech, Beijing) and were freshly used for flow cytometry analysis.

### Flow cytometry

Surface and intracellular staining for flow cytometry analysis were performed as described previously^49^. The antibodies used for surface and intracellular staining are listed in Table S3. For surface staining, cells were incubated with relevant fluorochrome-labeled antibodies for 20 min at 4°C in the dark. For intracellular cytokine staining, cells were fixed and permeabilized using the Intracellular Fixation & Permeabilization Buffer Set (Invitrogen, USA) and stained with FITC -anti-IFN-γ, PE-anti-IL-2, or APC-anti-TNF-α (eBioscience, San Diego, USA). Freshly isolated cells were used for all assays, and approximately 20 000-40 000 T cells were acquired for each sample using a BD FACS Canto II flow cytometer. Data analysis was performed using Flow Jo software V10.0.7 (Tree Star, Ashland, OR, USA). Cell debris and dead cells were excluded from the analysis based on scatter signals and Fixable Viability Dye eFluor 506 (Thermo Fisher Scientific).

### Analysis of the HBV-specific CD8^+^ T cell response in patients

The HBV-specific CD8^+^ T cells were detected after antigen-specific expansion as previously described^50^. Briefly, PBMCs were resuspended in complete medium (RPMI 1640 containing 10% fetal calf serum, 100U/ml penicillin, 100μg/ml streptomycin, and 100μM HEPES (4-[2-hydroxyethyl]-1-piperazine ethanesulfonic acid buffer) and stimulated with overlapping peptide pools covering the entire sequences of HBcAg or HBsAg (genotype B and C, GeneBank accession number: AF121243 and AF 112063), anti-CD28/CD49d (0.5μg/ml; BD Biosciences), and recombinant interleukin-2 (20U/ml; Hoffmann-La Roche). Fresh medium containing IL-2 was added twice per week. On day 10, the cells were tested for the expression of IFN-γ, TNF-α and IL-2 after re-stimulation with corresponding peptide pools by intracellular cytokine staining and subsequent flow cytometry analysis.

### Statistical analysis

Statistical analysis was performed using the SPSS statistical software package (version 22.0, SPSS Inc., Chicago, IL, USA). The Shapiro-Wilk method was used to test for normality. Parametric analysis methods were used when the data was normally distributed; otherwise, nonparametric tests were employed. Unpaired t test, paired t test, one-way ANOVA, Pearson product-moment correlation coefficient, log-rank test, and analysis of covariance (ANCOVA) were used where appropriate. The values of the selected parameters for predicting HBsAg loss in cHBV patients were assessed by receiver operating characteristic (ROC) and area under the ROC curve (AUC). All reported P values were two-sided, and a P value less than 0.05 was considered statistically significant.

## Acknowledgments

This work is supported by the National Natural Science Foundation of China (81861138044, 91742114, 81271884 and 81461130019), and the National Scientific and Technological Major Project of China (2017ZX10202203, 2018ZX10723203, 2018ZX10302206, 2017ZX10202201, 2017ZX10202202 and 2017YFC0908100), the Integrated Innovative Team for Major Human Diseases Program of Tongji Medical College and the “Double-First Class” Project for the International Cooperation Center on Infection and Immunity, HUST.

## Authors contributions

Conceived and designed the experiments: SEX, DZ, JL, XZ. Performed the experiments: SEX, DZ, BYL, MYL, WP, JYH. Analyzed and interpreted the data: SEX, DZ, JL, XZ. Contributed reagents/materials/analysis tools: HW, MJL, DLY, JL, XZ. Drafted the manuscript: SEX, MJL, KS, UD, DL, XZ, JL.

## Notes

### Competing Interest Statement

The authors have declared no competing interest.

