## supplementary tables and figures for "Longitudinal characterization of phenotypic profile of T cells in chronic hepatitis B identifies immune markers predicting HBsAg loss"

#### Table S1. Demographics and baseline characteristics of study subjects.

| | Healthy controls | HBsAg retained | HBsAg loss |
|---|---|---|---|
| **Number of subjects** | 24 | 141 | 31 |
| **Age, mean (range)** | 25.54±2.65 (21-34) | 33.33±7.75 (20-55) | 36.71±8.73 (18-55) |
| **Gender, male (%)** | 33.33% (8) | 76.60% (108) | 83.87% (26) |
| **ALT U/L, mean (range)** | n.a. | 32.86±39.13 (7-391) | 32.74±24.12 (9-122) |
| **HBeAg status, pos (%)** | n.a | 46.81% (66) | n.a. |
| **HBeAg, $\log_{10}$ index/ml, mean (range)** | n.a. | 2.07±1.01(0.45-4.62) | n.a. |
| **HBsAg, $\log_{10}$ IU/ml, mean (range)** | n.a. | 2.19±1.32(-0.92-4.99) | n.a. |
| **HBV DNA, neg (%)** | n.a. | 82.98% (117) | n.a. |
| **HBV DNA, $\log_{10}$ IU/ml, mean (range)** | n.a | 4.29±1.75(2.06-8.14) | n.a |
| **HBsAb status, pos (%)** | n.a | n.a | 51.61% (16) |
| **Treatment, IFN (%)** | n.a | 1.42% (2) | 16.13% (5) |
| **Treatment, NUCs+IFN (%)** | n.a | 8.51% (12) | 41.94% (13) |
| **Treatment, NUCs (%)** | n.a | 71.63% (101) | 19.35% (6) |
| **Without treatment (%)** | n.a | 18.44% (26) | 22.58% (7) |

Abbreviations: n.a., not applicable.

**Table S2. The details and characteristics of each patient enrolled in follow-up cohorts.**

Group A: sAg-ND, n=7

| Patient | Weeks | HBsAg IU/ml | HBsAb mIU/ml | HBeAg index/ml | HBeAb index/ml | HBcAb index/ml | HBV DNA | ALT | AST | Treatment | Immune test |
|---|---|---|---|---|---|---|---|---|---|---|---|
| #1 | 0 | 332.84 | 0.00 | 0.41 | >400.00 | 319.00 | | 19.00 | 16.00 | NUA | Y |
| | 13 | 352.69 | 0.37 | 0.56 | >400.00 | 323.10 | <100.00 | | | NUA | NA |
| | 31 | 352.92 | 0.67 | 0.32 | >400.00 | 318.80 | <100.00 | | | NUA | NA |
| | 38 | 379.53 | 1.08 | 0.25 | >400.00 | 443.30 | <100.00 | 18.00 | 18.00 | NUA | Y |
| | 60 | 378.47 | 1.22 | 0.00 | >400.00 | >500.00 | | 25.00 | 19.00 | NUA | Y |
| #2 | 0 | 364.03 | 1.34 | 1.02 | >400.00 | 403.00 | | | | NUA | Y |
| | 13 | 510.18 | 0.72 | 0.19 | >400.00 | 395.80 | | 10.00 | 18.00 | NUA | Y |
| | 26 | 361.71 | 0.22 | 1.31 | >400.00 | 446.50 | | 17.00 | 20.00 | NUA | Y |
| | 39 | 358.42 | 1.95 | 0.75 | >400.00 | >500.00 | | 16.00 | 20.00 | NUA | NA |
| | 55 | 318.48 | 0.10 | 0.48 | >400.00 | >500.00 | | 17.00 | 21.00 | NUA | Y |
| #3 | 0 | 11.80 | 0.33 | 0.13 | >400.00 | 392.40 | | 9.00 | 26.00 | NUA | Y |
| | 4 | 10.35 | 1.05 | 0.33 | >400.00 | 399.10 | | 13.00 | 21.00 | NUA | Y |
| | 12 | | | | | | | 11.00 | 23.00 | NUA | NA |
| | 13 | 10.83 | 0.00 | 0.49 | >400.00 | 403.20 | | | | NUA | Y |
| | 23 | 15.91 | 1.21 | 0.27 | >400.00 | 479.20 | <100.00 | 11.00 | 21.00 | NUA | NA |
| | 28 | | | | | | | 12.00 | 20.00 | NUA | Y |
| | 38 | 16.56 | 0.58 | 0.32 | >400.00 | >500.00 | | 8.00 | 17.00 | NUA | NA |
| | 52 | 24.84 | 0.35 | | | | | 6.00 | 14.00 | NUA | NA |
| #4 | 0 | 8.06 | 4.32 | 1.36 | 72.58 | 422.00 | | 11.00 | 15.00 | NUA | Y |
| | 9 | 6.76 | 3.26 | 0.42 | 56.38 | 471.50 | | 9.00 | 13.00 | NUA | NA |
| | 22 | 10.80 | 2.79 | 0.38 | 84.02 | 485.90 | | 10.00 | 14.00 | NUA | NA |
| | 29 | 3.76 | 5.31 | 0.17 | 109.50 | 487.60 | | 8.00 | 16.00 | NUA | Y |
| | 47 | 7.00 | 3.58 | 0.24 | 139.30 | >500.00 | <100.00 | 8.00 | 14.00 | NUA | Y |
| #5 | 0 | 0.43 | 0.61 | 1.96 | >400.00 | 400.30 | <100.00 | 48.00 | 41.00 | NUA | Y |
| | 5 | 0.85 | 0.76 | 0.11 | >400.00 | 394.40 | | 31.00 | 33.00 | NUA | Y |
| | 9 | 1.42 | 1.08 | 0.34 | >400.00 | 402.00 | | 60.00 | 30.00 | NUA | Y |
| | 13 | | | | | | | 110.00 | 45.00 | NUA | NA |
| | 33 | 1.92 | 1.33 | 0.41 | >400.00 | >500.00 | | 64.00 | 32.00 | NUA | Y |
| #6 | 0 | 425.37 | 1.35 | 35.55 | 12.36 | 370.10 | | 28.00 | 21.00 | NUA | Y |
| | 8 | 354.19 | 1.02 | 62.37 | 15.25 | 389.16 | | 31.00 | 27.00 | NUA | NA |
| | 21 | 521.07 | 1.12 | 67.52 | 3.10 | 354.20 | | | | NUA | NA |
| | 29 | 1000.00 | 0.98 | 53.27 | 0.68 | >500.00 | | | | NUA | Y |
| | 43 | 510.90 | | 85.10 | | | | 32.00 | 21.00 | NUA | NA |
| | 54 | 441.93 | 0.91 | 80.62 | 0.00 | >500.00 | | 29.00 | 24.00 | NUA | Y |
| #7 | 0 | 1.24 | 4.42 | 1.32 | 132.67 | 341.07 | <100.00 | | | NUA | Y |
| | 38 | 0.96 | 0.00 | 0.84 | 135.30 | 343.20 | | 7.00 | 15.00 | NUA | Y |
| | 60 | 1.01 | 0.00 | 0.32 | 105.70 | 334.60 | <100.00 | | | NUA | NA |

### Group B: sAg-RD50, n=5

| Patient | Weeks | HBsAg IU/ml | HBsAb mIU/ml | HBeAg index/ml | HBeAb index/ml | HBcAb index/ml | HBV DNA | ALT | AST | Treatment | Immune test |
|---|---|---|---|---|---|---|---|---|---|---|---|
| #8 | 0 | 466.63 | 1.00 | 0.30 | >400.00 | 417.40 | | 248.00 | 124.00 | NUA+IFN | Y |
| | 1 | | | | | | | 251.00 | 130.00 | NUA+IFN | NA |
| | 15 | 4.52 | 1.02 | 0.30 | >400.00 | 424.00 | | | | NUA+IFN | NA |
| | 17 | | | | | | | 116.00 | 76.00 | NUA+IFN | Y |
| | 23 | 0.33 | 2.49 | 0.15 | >400.00 | 403.80 | | | | NUA+IFN | NA |
| | 28 | 0.13 | 2.08 | 0.21 | >400.00 | 378.00 | | 47.00 | 38.00 | NUA+IFN | Y |
| | 35 | 0.58 | 1.39 | 0.28 | >400.00 | 472.30 | | 26.00 | 29.00 | NUA+IFN | NA |
| | 43 | 3.75 | 2.48 | 0.26 | >400.00 | 484.10 | | 26.00 | 29.00 | NUA+IFN | NA |
| | 44 | | | | | | | 26.00 | 29.00 | NUA+IFN | Y |
| | 54 | 0.11 | 1.78 | 0.00 | >400.00 | 496.40 | | 65.00 | 46.00 | NUA+IFN | Y |
| | 59 | 3.25 | 1.96 | 0.00 | >400.00 | >500 | | 23.00 | 24.00 | NUA+IFN | NA |
| #9 | 0 | 142.79 | 1.39 | 0.03 | >400.00 | >500.00 | <100.00 | 22.00 | 20.00 | NUA | Y |
| | 18 | 60.40 | 2.45 | 0.00 | >400.00 | >500.00 | <100.00 | 21.00 | 21.00 | NUA+IFN | Y |
| | 25 | 13.77 | 1.11 | 0.00 | >400.00 | >500.00 | <100.00 | 53.00 | 59.00 | NUA+IFN | NA |
| | 34 | 2.22 | 1.53 | 0.09 | >400.00 | >500.00 | <100.00 | 29.00 | 27.00 | NUA+IFN | Y |
| | 51 | 2.62 | 1.23 | 0.38 | >400.00 | >500.00 | <100.00 | 31.00 | 29.00 | NUA+IFN | NA |
| #10 | 0 | 91.08 | 0.42 | 5.59 | 137.10 | 481.60 | <100.00 | 17.00 | 17.00 | NUA | Y |
| | 5 | | | | | | | 77.00 | 43.00 | NUA+IFN | Y |
| | 17 | 10.85 | 2.47 | 0.23 | >400.00 | 497.60 | | | | NUA+IFN | NA |
| | 21 | 5.68 | 0.98 | 1.07 | >400.00 | 495.70 | | 98.00 | 53.00 | NUA+IFN | Y |
| | 25 | 3.94 | 1.48 | 0.00 | >400.00 | >500.00 | | 87.00 | 46.00 | NUA+IFN | Y |
| | 30 | 2.27 | 1.07 | 0.00 | >400.00 | >500.00 | | 49.00 | 30.00 | NUA+IFN | NA |
| | 34 | 0.82 | 3.56 | 0.11 | >400.00 | >500.00 | | 98.00 | 47.00 | NUA+IFN | NA |
| | 38 | 0.14 | 5.11 | 0.29 | >400.00 | >500.00 | | 68.00 | 39.00 | NUA+IFN | Y |
| | 43 | 0.15 | 7.80 | 0.21 | >400.00 | >500.00 | | 32.00 | 26.00 | NUA+IFN | NA |
| | 47 | | | | | | | | | NUA+IFN | NA |
| | 52 | 0.11 | 5.25 | 0.29 | >400.00 | >500.00 | <100.00 | 12.00 | 15.00 | NUA+IFN | Y |
| #11 | 0 | 61.89 | 0.25 | 3.40 | 26.14 | 319.00 | | 18.00 | 16.00 | NUA | Y |
| | 9 | 56.20 | 0.63 | 7.12 | 27.09 | 290.60 | <100.00 | | | NUA | Y |
| | 16 | 84.60 | 0.12 | 6.93 | 27.86 | 275.70 | | 33.00 | 23.00 | NUA | NA |
| | 24 | | | | | | | 36.00 | 30.00 | NUA | NA |
| | 27 | 83.81 | 0.00 | 27.40 | 27.12 | 293.40 | | | | NUA | NA |
| | 30 | | | | | | | 25.00 | 24.00 | NUA+IFN | NA |
| | 36 | 85.86 | 0.63 | 18.37 | 29.25 | 251.50 | | 32.00 | 24.00 | NUA+IFN | Y |
| | 44 | 3.29 | 0.54 | 15.03 | 11.74 | 225.70 | | 62.00 | 39.00 | NUA+IFN | NA |
| | 53 | 3.59 | 0.00 | 3.11 | 49.64 | 421.10 | | 44.00 | 29.00 | NUA+IFN | NA |
| #12 | 0 | 59.47 | 1.57 | 0.19 | 41.91 | 466.50 | | 29.00 | 21.00 | NUA | Y |
| | 13 | 72.87 | 0.73 | 0.00 | 38.91 | >500.00 | | 41.00 | 26.00 | NUA+IFN | Y |
| | 22 | 30.77 | 0.58 | 0.00 | 54.82 | >500.00 | | 60.00 | 39.00 | NUA+IFN | Y |
| | 26 | 9.30 | 1.03 | 0.00 | 43.16 | >500.00 | | 68.00 | 45.00 | NUA+IFN | NA |

| | | | | | | | | | | | |
|---|---|---|---|---|---|---|---|---|---|---|---|
| | 31 | 3.69 | 1.28 | 0.00 | 51.88 | >500.00 | | 198.00 | 100.00 | NUA+IFN | NA |
| | 32 | 1.80 | 2.05 | 0.22 | 46.14 | >500.00 | | 167.00 | 72.00 | NUA+IFN | Y |
| | 34 | 1.23 | 1.13 | 0.10 | 43.59 | >500.00 | | 165.00 | 188.00 | NUA+IFN | NA |
| | 38 | 0.56 | 3.49 | 0.27 | 51.92 | >500.00 | <100.00 | 65.00 | 40.00 | NUA+IFN | Y |
| | 43 | 0.31 | 4.45 | 0.30 | 42.38 | >500.00 | | 65.00 | 43.00 | NUA+IFN | NA |
| | 48 | | | | | | | | | NUA+IFN | NA |
| | 52 | 0.22 | 5.28 | 0.45 | 32.22 | >500.00 | <100.00 | 55.00 | 40.00 | NUA+IFN | Y |

#### Group C: sAg-L, n=6

| **Patient** | **Weeks** | **HBsAg IU/ml** | **HBsAb mIU/ml** | **HBeAg index/ml** | **HBeAb index/ml** | **HBcAb index/ml** | **HBV DNA** | **ALT** | **AST** | **Treatment** | **Immune test** |
|---|---|---|---|---|---|---|---|---|---|---|---|
| #13 | 0 | 500.15 | 0.61 | 0.28 | >400.00 | 391.80 | 7.18E+05 | 183.00 | 49.00 | NUA+IFN | Y |
| | 17 | | | | | | | 32.00 | 25.00 | NUA+IFN | NA |
| | 26 | 103.09 | 2.94 | 0.53 | >400.00 | 493.20 | 2.14E+03 | 26.00 | 24.00 | NUA+IFN | NA |
| | 30 | | | | | | | 68.00 | 38.00 | NUA+IFN | NA |
| | 34 | 111.42 | 1.37 | 0.44 | >400.00 | >500.00 | <100.00 | 66.00 | 35.00 | NUA+IFN | NA |
| | 38 | | | | | | | 62.00 | 38.00 | NUA+IFN | NA |
| | 43 | 2.57 | 10.10 | 0.34 | >400.00 | >500.00 | | 46.00 | 35.00 | NUA+IFN | Y |
| | 46 | | | | | | | 122.00 | 58.00 | NUA+IFN | NA |
| | 47 | 1.91 | 6.74 | | | | | 57.00 | 45.00 | NUA+IFN | NA |
| | 51 | 0.02 | 30.46 | 0.50 | >400.00 | 478.40 | | 33.00 | 36.00 | NUA+IFN | Y |
| | 56 | 0.03 | 40.07 | 0.41 | >400.00 | 482.60 | | 79.00 | 47.00 | NUA+IFN | NA |
| | 60 | 0.07 | 34.07 | 0.60 | >400.00 | >500 | <100 | | | | NA |
| #14 | 0 | 6.50 | 0.70 | 0.19 | 28.36 | 385.96 | <100.00 | 10.00 | 17.00 | IFN | Y |
| | 4 | 6.48 | 1.16 | 0.29 | 26.67 | 380.66 | | | | IFN | NA |
| | 13 | 1.52 | 1.07 | 0.07 | 25.48 | 401.20 | | | | IFN | NA |
| | 16 | | | | | | | | | IFN | Y |
| | 20 | 0.58 | 0.68 | 1.15 | 25.49 | 423.00 | | | | IFN | Y |
| | 23 | 0.57 | 0.48 | 1.95 | 39.08 | 385.50 | | | | IFN | NA |
| | 27 | 0.31 | 1.60 | 0.26 | 31.89 | 397.30 | | | | IFN | Y |
| | 31 | 0.19 | 1.93 | 0.53 | 30.72 | 386.30 | | 15.00 | 19.00 | IFN | Y |
| | 36 | 0.10 | 0.00 | 0.54 | 37.83 | 408.60 | | 14.00 | 23.00 | IFN | Y |
| | 40 | 0.05 | 1.05 | 0.59 | 34.06 | 416.10 | | | | IFN | Y |
| | 53 | 0.05 | 1.98 | 0.72 | 34.54 | >500.00 | | 13.00 | 22.00 | IFN | NA |
| | 57 | 0.04 | 0.94 | 0.39 | 23.59 | >500.00 | | 14.00 | 21.00 | IFN | Y |
| #15 | 0 | 6.84 | 1.78 | 0.36 | >400.00 | 409.50 | <100.00 | 18.00 | 18.00 | NUA+IFN | Y |
| | 1 | | | | | | | 12.00 | 19.00 | NUA+IFN | NA |

| | | | | | | | | | | | |
|---|---|---|---|---|---|---|---|---|---|---|---|
| | 3 | | | | | | | 32.00 | 31.00 | NUA+IFN | NA |
| | 4 | | | | | | | 49.00 | 42.00 | NUA+IFN | NA |
| | 6 | 11.09 | 0.64 | 0.22 | >400.00 | 490.30 | | 32.00 | 26.00 | NUA+IFN | NA |
| | 8 | 4.78 | 0.96 | 0.40 | >400.00 | >500.00 | <100.00 | 29.00 | 30.00 | NUA+IFN | NA |
| | 10 | | | | | | | 28.00 | 29.00 | NUA+IFN | NA |
| | 12 | 0.57 | 0.66 | 0.42 | >400.00 | 461.40 | | 48.00 | 39.00 | NUA+IFN | Y |
| | 13 | 0.80 | | | | | | 32.00 | 29.00 | NUA+IFN | NA |
| | 15 | 0.17 | 1.91 | 0.20 | >400.00 | >500.00 | | 36.00 | 34.00 | NUA+IFN | Y |
| | 17 | 0.11 | 0.77 | 0.27 | >400.00 | 453.10 | | | | NUA+IFN | NA |
| | 19 | 0.11 | 0.91 | 0.40 | >400.00 | >500.00 | | | | NUA+IFN | NA |
| | 25 | 0.06 | 0.92 | 0.28 | >400.00 | >500.00 | | 20.00 | 19.00 | NUA+IFN | Y |
| | 28 | 0.26 | 4.20 | 0.27 | >400.00 | >500.00 | | 13.00 | 22.00 | NUA+IFN | Y |
| | 31 | 0.05 | 1.05 | 0.07 | >400.00 | >500.00 | | 18.00 | 22.00 | NUA+IFN | Y |
| | 35 | 0.10 | 2.41 | 0.07 | >400.00 | 496.10 | | 20.00 | 19.00 | NUA+IFN | NA |
| | 53 | 0.36 | 1.55 | 0.03 | >400.00 | 497.70 | | 16.00 | 17.00 | NUA+IFN | Y |
| | 57 | 0.85 | 1.39 | 0.19 | >400.00 | 496.20 | | 14.00 | 16.00 | NUA+IFN | NA |
| #16 | 0 | 5.54 | | | | | | 36.00 | 25.00 | NUA+IFN | Y |
| | 2 | 13.54 | 0.98 | 0.16 | >400.00 | 494.00 | | 50.00 | 38.00 | NUA+IFN | Y |
| | 6 | 0.41 | 1.46 | 0.44 | >400.00 | 479.50 | | 65.00 | 40.00 | NUA+IFN | Y |
| | 11 | 0.09 | 1.50 | 0.30 | >400.00 | >500.00 | | 32.00 | 28.00 | NUA+IFN | Y |
| | 16 | 0.13 | 0.31 | 0.05 | 361.80 | 499.00 | | 25.00 | 24.00 | NUA+IFN | Y |
| | 21 | 0.07 | 3.33 | 0.00 | >400.00 | >500.00 | | 21.00 | 22.00 | NUA+IFN | NA |
| | 24 | 0.02 | 3.07 | 0.00 | >400.00 | >500.00 | | | | NUA+IFN | NA |
| | 30 | 0.02 | 2.43 | 0.00 | 341.80 | >500.00 | | 17.00 | 19.00 | NUA+IFN | NA |
| | 35 | 0.08 | 2.49 | 0.20 | 386.10 | >500.00 | | 25.00 | 21.00 | NUA+IFN | Y |
| | 41 | 0.05 | 2.48 | 0.27 | >400.00 | >500.00 | | 19.00 | 17.00 | NUA+IFN | NA |
| | 45 | 0.04 | 1.73 | 0.22 | 379.00 | >500.00 | | 26.00 | 18.00 | NUA+IFN | NA |
| #17 | 0 | 0.67 | 3.33 | 0.29 | >400.00 | 332.70 | | | | NUA+IFN | Y |
| | 10 | 1.07 | 9.61 | 1.51 | >400.00 | 318.54 | | | | NUA+IFN | NA |
| | 20 | 0.15 | 14.41 | 0.86 | >400.00 | 362.10 | | | | NUA+IFN | NA |
| | 29 | 0.09 | 10.86 | 0.02 | >400.00 | 391.30 | | | | NUA+IFN | Y |
| | 34 | 0.14 | 6.44 | 0.00 | >400.00 | 403.20 | | | | NUA+IFN | Y |
| | 43 | 0.05 | 26.09 | 0.18 | >400.00 | 362.70 | | 29.00 | 30.00 | NUA+IFN | Y |
| | 52 | 0.01 | 73.50 | 0.34 | >400.00 | 389.60 | | | | NUA+IFN | Y |

| | | | | | | | | | | | |
|---|---|---|---|---|---|---|---|---|---|---|---|
| | 59 | 0.00 | 40.10 | 0.58 | >400.00 | 375.30 | | 20.00 | 27.00 | NUA+IFN | Y |
| #18 | 0 | 0.13 | 1.86 | 3.02 | 36.41 | 412.90 | | 19.00 | 23.00 | NUA+IFN | Y |
| | 3 | | | | | | | 21.00 | 24.00 | NUA+IFN | NA |
| | 8 | 0.11 | 3.94 | 2.60 | 18.59 | 443.20 | <100.00 | 17.00 | 22.00 | NUA+IFN | NA |
| | 13 | 0.07 | 6.34 | 2.12 | 353.00 | 441.30 | | 23.00 | 27.00 | NUA+IFN | NA |
| | 17 | 0.13 | 3.15 | 2.55 | 17.70 | 447.50 | <100.00 | 12.00 | 19.00 | NUA+IFN | NA |
| | 21 | 0.10 | 3.60 | 2.16 | 15.60 | 465.40 | | 16.00 | 23.00 | NUA+IFN | NA |
| | 26 | 0.08 | 6.13 | 2.21 | 14.91 | 450.80 | <100.00 | 13.00 | 22.00 | NUA+IFN | Y |
| | 30 | 0.03 | 11.19 | 2.34 | 12.07 | 454.30 | | 12.00 | 20.00 | NUA+IFN | Y |
| | 39 | 0.01 | 12.95 | 1.41 | 5.74 | >500.00 | <100.00 | 15.00 | 23.00 | NUA+IFN | NA |
| | 42 | | | | | | | 13.00 | 21.00 | NUA+IFN | Y |
| | 47 | 0.04 | 4.75 | 1.38 | 2.04 | 490.60 | <100.00 | 13.00 | 21.00 | NUA+IFN | NA |
| | 51 | 0.03 | 4.41 | 1.78 | 46.16 | >500.00 | | 14.00 | 21.00 | NUA+IFN | NA |
| | 56 | 0.03 | 7.00 | 1.85 | 40.24 | >500.00 | | 15.00 | 21.00 | NUA+IFN | NA |
| | 60 | 0.04 | 5.43 | 2.62 | 35.04 | >500 | <100.00 | 25.00 | 21.00 | NUA+IFN | NA |

ALT: alanine aminotransferase

AST: aspartate aminotransferase

NA: not available

#### Table S3. Antibodies used for flow cytometry analysis

| Antibody | Colour | Clone | Company |
|---|---|---|---|
| **CD3** | APC-eFluor® 780 | SK7 | eBioscience |
| **CD4** | PerCP-Cyanine5.5 | OKT4 | eBioscience |
| **CD8** | PE-Cyanine7 | SK1 | eBioscience |
| **CD8** | FITC | HIT8a | eBioscience |
| **CD19** | PE | HIB19 | eBioscience |
| **CD11c** | APC | 3.9 | eBioscience |
| **CD11c** | APC-eFluor® 780 | 3.9 | eBioscience |
| **CD14** | eFluor® 450 | 61D3 | eBioscience |
| **CD43** | FITC | eBio84-3C1 | eBioscience |
| **Granzyme B*** | PE | GB11 | eBioscience |
| **CD62L** | APC | DREG-56 | eBioscience |
| **CD69** | eFluor® 450 | FN50 | eBioscience |
| **CD107a** | PE | eBioH4A3 | eBioscience |
| **HLA-DR** | PE-Cyanine7 | L243 | eBioscience |
| **CD95** | APC | DX2 | eBioscience |
| **CD40L** | eFluor® 450 | 24-31 | eBioscience |
| **Tim-3** | FITC | F38-2E2 | eBioscience |
| **CTLA-4** | PE | eBio20A | eBioscience |
| **PD-1** | APC | eBioJ105 | eBioscience |
| **CD25** | eFluor® 450 | BC96 | eBioscience |
| **CD86** | Alexa Fluor® 488 | IT2.2 | eBioscience |
| **PD-L1** | PerCP-eFluor™ 710 | MIH1 | eBioscience |
| **CD80** | PE-Cy™7 | L307.4 | BD Bioscience |
| **CD72** | eFluor® 660 | J4-117 | eBioscience |
| **IFN-γ** | FITC | 4S.B3 | eBioscience |
| **TNF-a** | APC | MAb11 | eBioscience |
| **IL-2** | PE | MQ1-17H12 | eBioscience |
| **CD3** | Functional Grade | OKT3 | eBioscience |
| **CD28** | Functional Grade | CD28.2 | eBioscience |

*Intracellular antibodies

**Table S4. Baseline characteristics of sAg-ND and sAg-RD patients.**

| | **sAg-ND** | **sAg-RD30** | **sAg-RD50** |
|---|---|---|---|
| Number of subjects | 30 | 19 | 8 |
| ALT U/L, mean (range) | 26.87±24.44 (8-116) | 48.95±42.74 (13-161) | 50.50±46.77 (17-159) |
| HBeAg status, pos (%) | 63.33% (19) | 57.90% (11) | 62.50% (5) |
| HBsAg, log $_{10}$ IU/ml, mean (range) | 2.58±0.73(0.51-4.21) | 2.48±0.91(0.17-4.25) | 1.78±0.87(0.17-2.90) |
| HBV DNA, neg (%) | 83.33% (25) | 94.74% (18) | 100% (8) |
| Treatment, NUA (%) | 100% (30) | 68.42% (13) | 37.50% (3) |
| Treatment, IFN+NUA (%) | 0.00% (0) | 31.58% (6) | 62.50% (5) |

**Table S5. The area under the curve (AUC) of predictive parameters.**

| | | | | 95% CI | |
|---|---|---|---|---|---|
| | Phenotypes | AUC | P value | Lower | Upper |
| HBsAg loss within 48 weeks | HLA-DR$^+$CD4+ T cells | 0.773 | 0.031 | 0.607 | 0.939 |
| | CTLA-4$^+$CD4+ T cells | 0.785 | 0.024 | 0.608 | 0.963 |
| | HLA-DR$^+$CD8+ T cells | 0.807 | 0.015 | 0.637 | 0.977 |
| | TIM-3$^+$CD8+ T cells | 0.883 | 0.002 | 0.750 | 1.000 |
| | qHBsAg log IU/ml | 0.135 | 0.018 | 0.000 | 0.467 |

#### SUPPLEMENTARY FIGURE LEGENDS

**Figure S1. Analysis of different lymphocyte subsets in cHBV patients with HBsAg loss.** (A) Schematic representation of the study design. (B) Gating strategy of flow cytometry for different lymphocyte subsets. (C) Frequencies and numbers of T cells, B cells, monocytes, dendritic cells (DCs), CD4 and CD8 T cells were detected in healthy controls (HC, n=24), HBsAg retained patients (sAg-R, n=141) and HBsAg loss (sAg-L, n = 31) by flow cytometry. Error bars, mean ± s.e.m.; *P<0.05; **P<0.01; ***P<0.001; ****P<0.0001. ns, not significant.

**Figure S2. Characterization of T cell phenotype profiles in cHBV patients with HBsAg loss.** Expression of CD69, TIM-3, granzyme B, and CD107a on CD4 T cells, and CD69, CD25, CD40L, TIM-3, CTLA-4, Granzyme B, and CD107a on CD8 T cells of HC, sAg-R and sAg-L patients were analyzed by flow cytometry. Error bars, mean ± s.e.m.; *P<0.05; **P<0.01; ***P<0.001; ****P<0.0001. ns, not significant.

**Figure S3. Characterization of T cell phenotype profiles in sAg-L patients with seroconversion to anti-HBs.** (A) Immune marker expression on $CD4^+$ T cells of sAg-L patients with and without seroconversion to anti-HBs were analyzed by flow cytometry. (B) Immune marker expression on $CD8^+$ T cells of sAg-L patients with and without seroconversion to anti-HBs were analyzed by flow cytometry. Error bars, mean ± s.e.m.; ns, not significant.

**Figure S4. Characterization of T cell phenotype profiles in cHBV patients with**

**HBsAg rapid decrease.** Expression of CD69, PD-1, and granzyme B on CD4 T cells, and CD95, CD25, CD40L, PD-1, CTLA-4, and granzyme B on CD8 T cells were analyzed by flow cytometry and were compared between cHBV patients who experienced greater than 30% decrease of HBsAg levels compared to the baseline within 6 months (sAg-RD30, n=19), patients who experienced greater than 50% decrease of HBsAg levels compared to the baseline within 6 months (sAg-RD50, n=8), and patients with no decrease of serum HBsAg levels (sAg-ND, n=30). Error bars, mean ± s.e.m.; *p<0.05, **p<0.01, ****p<0.0001, ns-not significant (p>0.05).

**Figure S5. Pearson correlation analysis between T cell phenotypes and clinical parameters** (A) Pearson correlation analysis between the expression of corresponding T cell markers and the extent of HBsAg reduction was performed in CHB patients. (B) Pearson correlation analysis between the expression of corresponding T cell markers and serum HBsAg levels was performed in CHB patients. (C) Pearson correlation analysis between the expression of corresponding T cell markers and ALT levels was performed in CHB patients. (D) Pearson correlation analysis between the expression of corresponding T cell markers and HBV DNA levels was performed in CHB patients. Spearman's Rank correlation, ns-not significant (p>0.05).

**Figure S6. Analysis of HBcAg-specific T cell responses in sAg-L patients with seroconversion to anti-HBs.** (A) **The intensities of** HBcAg-specific CD4 and CD8 T cell responses were compared between sAg-L patients with and without seroconversion to anti-HBs. (B) Pearson correlation between CD4 T cell phenotypes and HBcAg-IFN-$\gamma^+$ CD4+ T cells (C) Pearson correlation between CD8 T cell phenotypes and HBcAg-IFN-$\gamma^+$ CD8+ T cells**.** Error bars, mean ± s.e.m.; ns-not significant ($p>0.05$).

**Figure S7. Characterization of HBsAg-specific T cell response in cHBV patients with HBsAg loss.** PBMCs from chronic hepatitis B patients were stimulated with overlapping peptide pools covering the entire sequences of HBsAg for 10 days. Cells were analyzed for IFN-γ, TNF-α, and IL-2 production by intracellular cytokine staining. (A) HBsAg -specific T cell responses were compared between sAg-R and sAg-L patients. (B) Pearson correlation analysis between expression of corresponding T cell markers and frequencies of IFN-γ-producing HBsAg-specific T cells was performed in CHB patients. Error bars, mean ± s.e.m.; ns-not significant ($p>0.05$).

**Figure S8. Characterization of T cell phenotype in cHBV patients received IFN-α treatment.** CD4 T cell (A)and CD8 T cell (B) phenotypes were compared between cHBV patients received NUC monotherapy (cHBV-NUC), IFN-α contained therapy (cHBV-IFN), and no therapy (CHB-NT). Error bars, mean ± s.e.m.; ns-not significant ($p>0.05$).

**Figure S9. Kinetic analysis of T cell phenotypes in cHBV patients during HBsAg rapid decrease and loss.** (A)The phenotypes of CD4 and CD8 T cells (B) were longitudinally analyzed during the observational period. Left: Data were pooled and compared at either baseline (green background) and follow-up period (pink background) between cHBV patients with no HBsAg decrease (sAg-ND), HBsAg rapid decrease (sAg-RD50), and HBsAg loss (sAg-L). Right: Kinetic changes of T cell phenotype and HBcAg-specific T cell responses were demonstrated at indicated time points for sAg-ND, sAg-RD50, and sAg-L patients. (C) Kinetic analysis of T cell phenotypes and HBcAg-specific T cell responses in individual cHBV patients. Data from 2 representative patients with no HBsAg reduction. (D) Kinetic analysis of T cell phenotypes and HBcAg-specific T cell responses in individual sAg-L patients. Data from 6 representative patients with HBsAg seroconversion.

### Figure S1

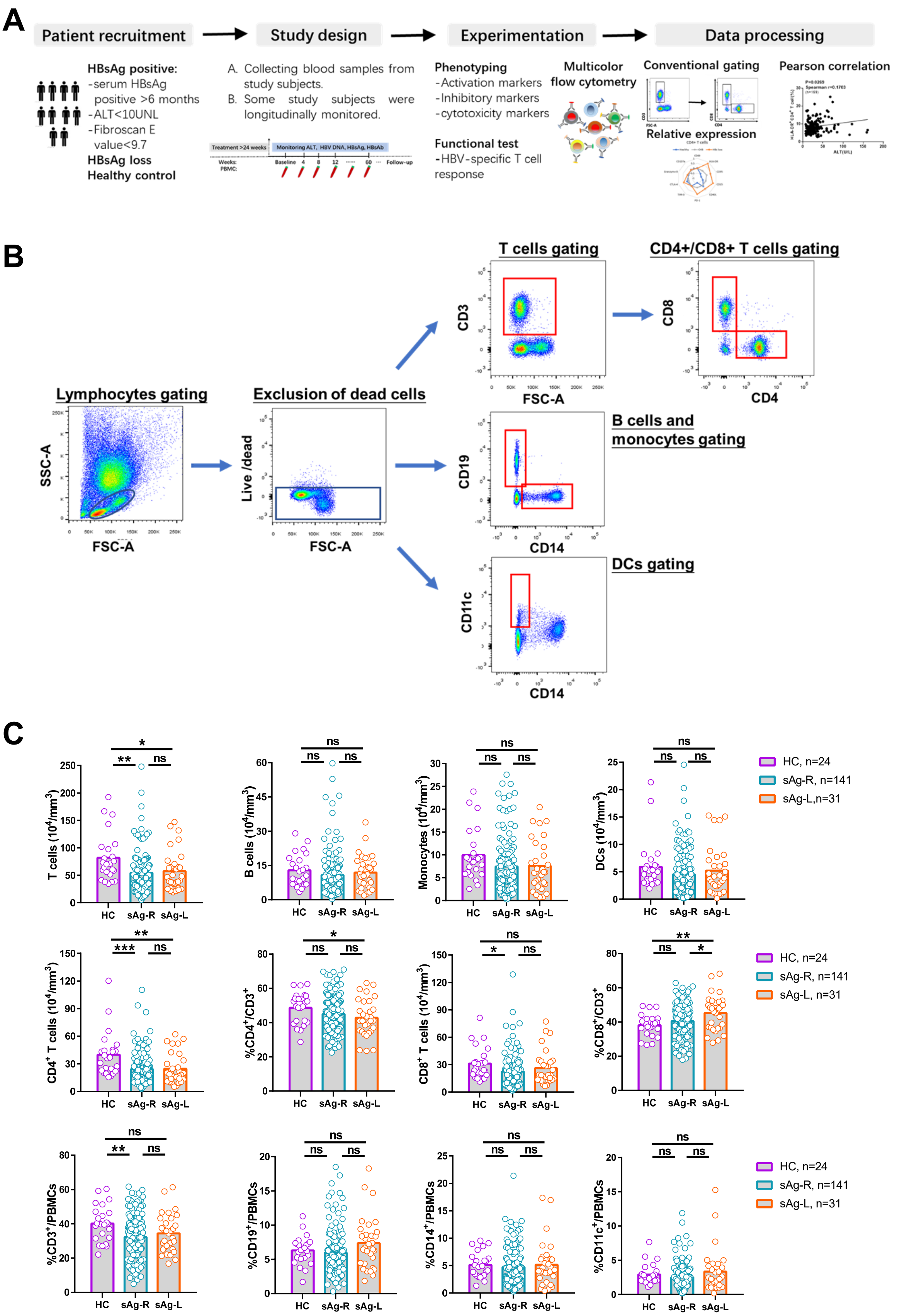
A
Patient recruitment
Study design
Experimentation
Data processing
HBsAg positive:
-serum HBsAg positive >6 months
-ALT<10UNL
-Fibroscan E value<9.7
HBsAg loss
Healthy control
A. Collecting blood samples from study subjects.
B. Some study subjects were longitudinally monitored.
Phenotyping
-Activation markers
-Inhibitory markers
-cytotoxicity markers
Multicolor flow cytometry
Functional test
-HBV-specific T cell response
Conventional gating
Relative expression
Pearson correlation
B
Lymphocytes gating
Exclusion of dead cells
T cells gating
CD4+/CD8+ T cells gating
B cells and monocytes gating
DCs gating
SSC-A
FSC-A
Live /dead
CD3
CD8
CD4
CD19
CD14
CD11c
C
HC, n=24
sAg-R, n=141
sAg-L,n=31
T cells (10^4/mm^3)
B cells (10^4/mm^3)
Monocytes (10^4/mm^3)
DCs (10^4/mm^3)
CD4+ T cells (10^4/mm^3)
%CD4+/CD3+
CD8+ T cells (10^4/mm^3)
%CD8+/CD3+
%CD3+/PBMCs
%CD19+/PBMCs
%CD14+/PBMCs
%CD11c+/PBMCs
HC
sAg-R
sAg-L

### Figure S2

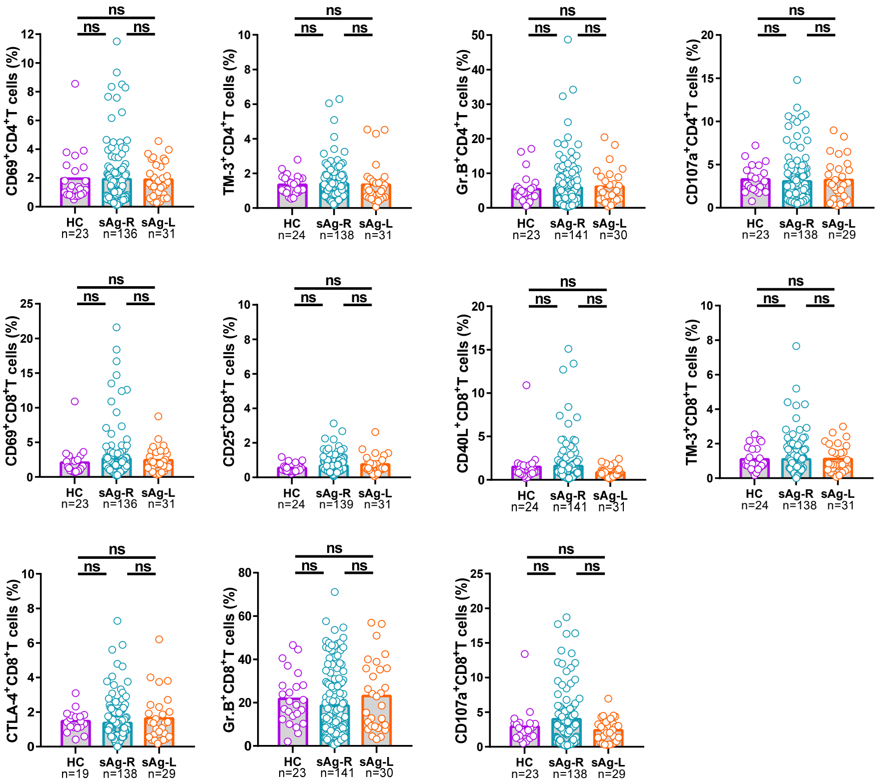

CD69+CD4+T cells (%)
TM-3+CD4+T cells (%)
Gr.B+CD4+T cells (%)
CD107a+CD4+T cells (%)
CD69+CD8+T cells (%)
CD25+CD8+T cells (%)
CD40L+CD8+T cells (%)
TM-3+CD8+T cells (%)
CTLA-4+CD8+T cells (%)
Gr.B+CD8+T cells (%)
CD107a+CD8+T cells (%)
ns
HC
sAg-R
sAg-L
n=23 n=136 n=31
n=24 n=138 n=31
n=23 n=141 n=30
n=23 n=138 n=29
n=23 n=136 n=31
n=24 n=139 n=31
n=24 n=141 n=31
n=24 n=138 n=31
n=19 n=138 n=29
n=23 n=141 n=30
n=23 n=138 n=29

### Figure S3

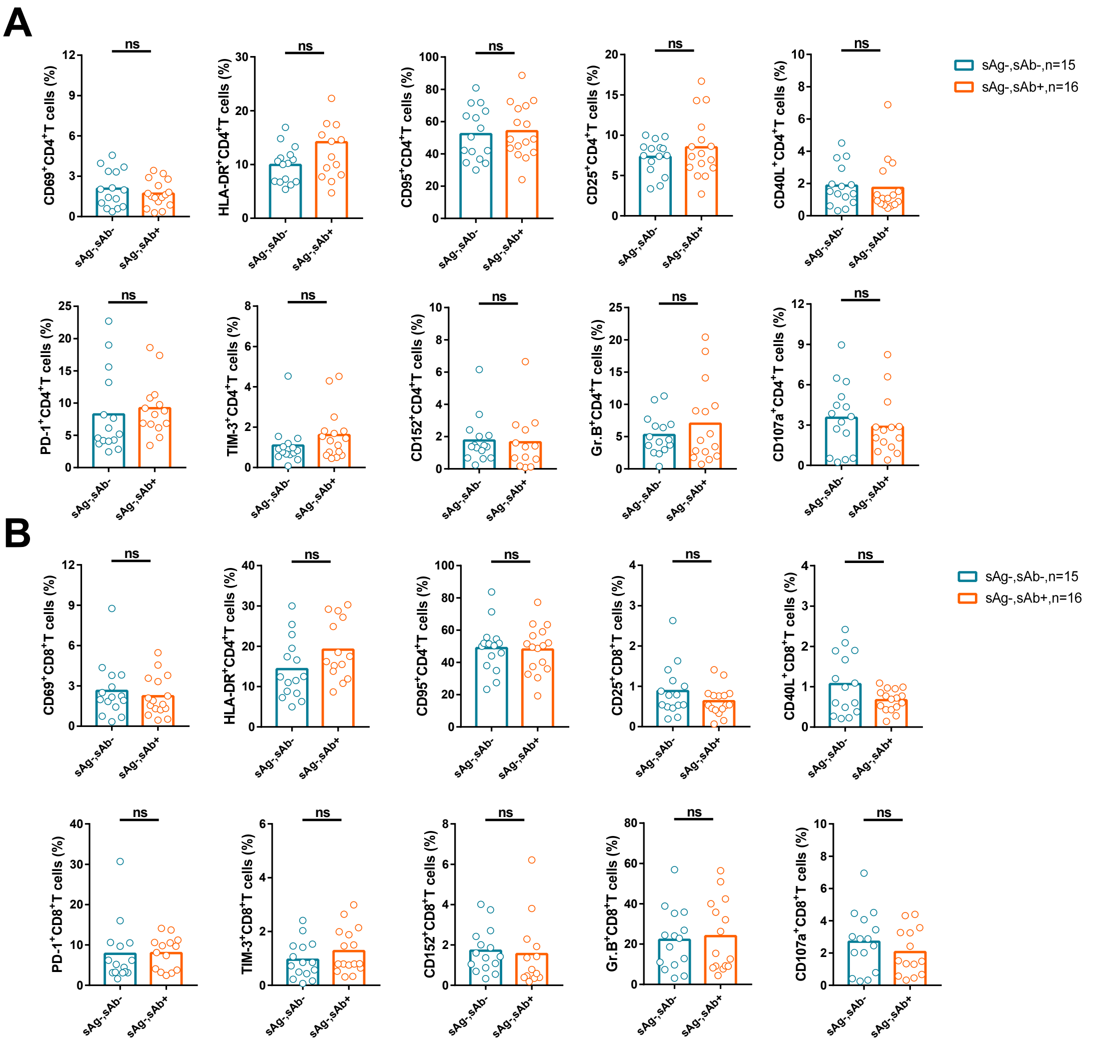
A
sAg-,sAb-,n=15
sAg-,sAb+,n=16
CD69+CD4+T cells (%)
HLA-DR+CD4+T cells (%)
CD95+CD4+T cells (%)
CD25+CD4+T cells (%)
CD40L+CD4+T cells (%)
PD-1+CD4+T cells (%)
TIM-3+CD4+T cells (%)
CD152+CD4+T cells (%)
Gr.B+CD4+T cells
CD107a+CD4+T cells (%)
ns
sAg-sAb-
sAg-sAb+
B
sAg-,sAb-,n=15
sAg-,sAb+,n=16
CD69+CD8+T cells (%)
HLA-DR+CD4+T cells (%)
CD95+CD4+T cells (%)
CD25+CD8+T cells (%)
CD40L+CD8+T cells (%)
PD-1+CD8+T cells (%)
TIM-3+CD8+T cells (%)
CD152+CD8+T cells (%)
Gr.B+CD8+T cells (%)
CD107a+CD8+T cells (%)
ns
sAg-sAb-
sAg-sAb+

### Figure S4

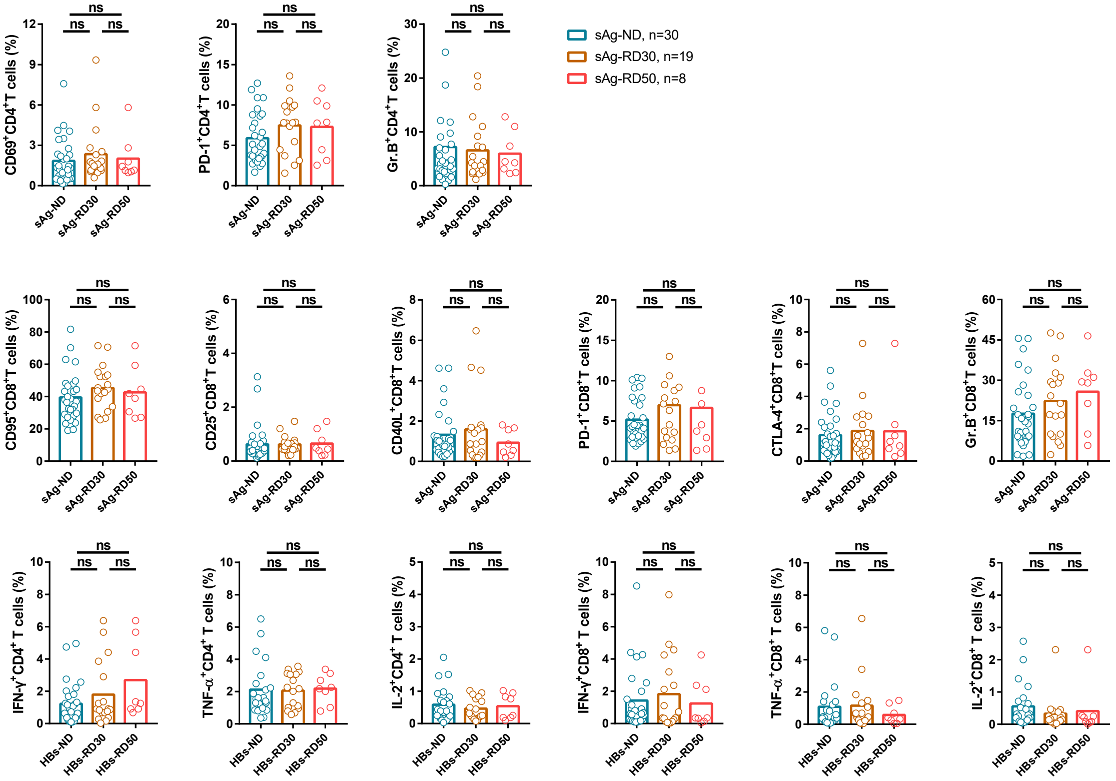
sAg-ND, n=30
sAg-RD30, n=19
sAg-RD50, n=8
ns
CD69+CD4+T cells (%)
PD-1+CD4+T cells (%)
Gr.B+CD4+T cells (%)
CD95+CD8+T cells (%)
CD25+CD8+T cells (%)
CD40L+CD8+T cells (%)
PD-1+CD8+T cells (%)
CTLA-4+CD8+T cells (%)
Gr.B+CD8+T cells (%)
IFN-γ+CD4+ T cells (%)
TNF-α+CD4+ T cells (%)
IL-2+CD4+ T cells (%)
IFN-γ+CD8+ T cells (%)
TNF-α+CD8+ T cells (%)
IL-2+CD8+ T cells (%)
sAg-ND
sAg-RD30
sAg-RD50
HBs-ND
HBs-RD30
HBs-RD50

### Figure S5

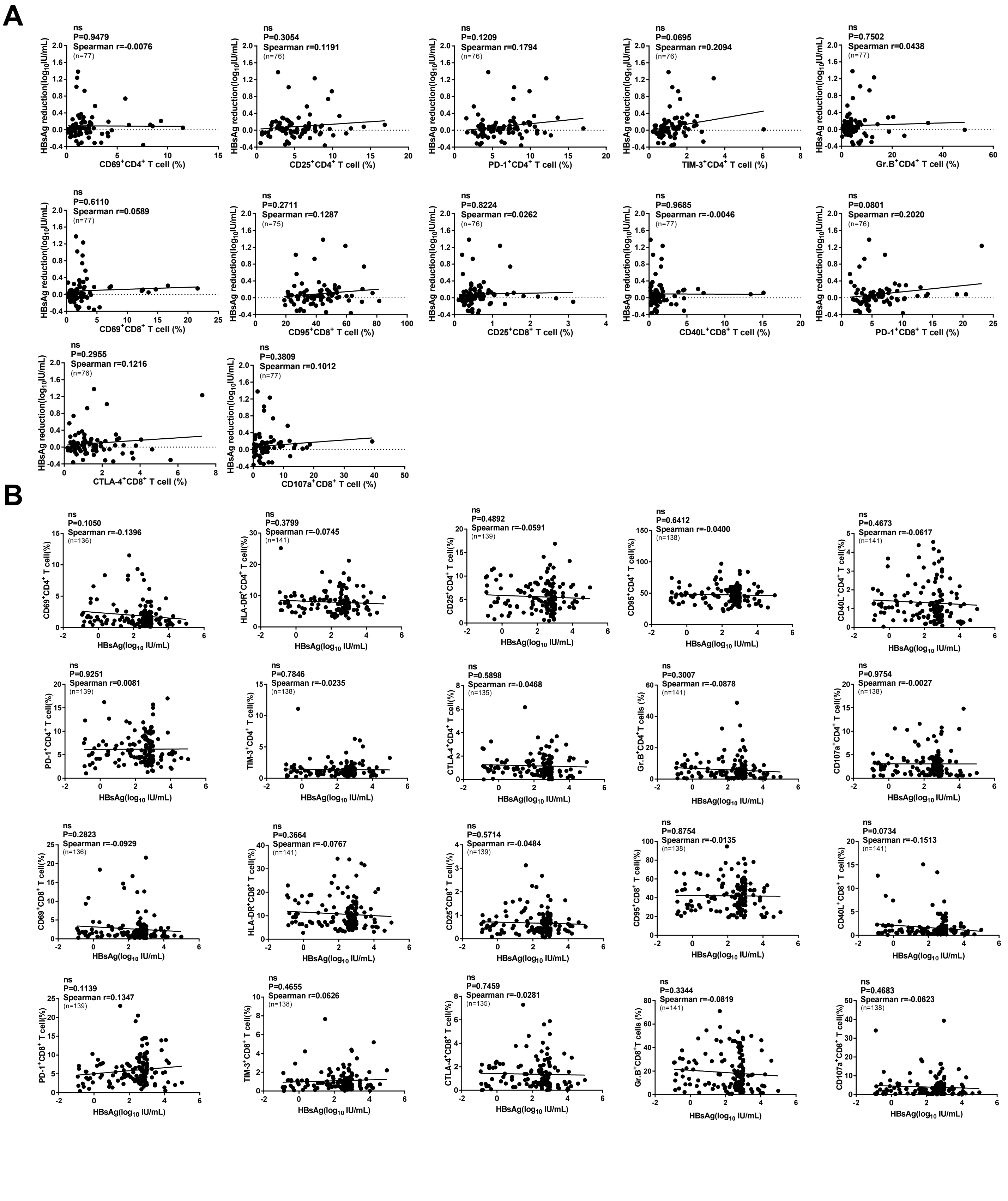
A
ns
P=0.9479
Spearman r=-0.0076
(n=77)
HBsAg reduction($log_{10}$IU/mL)
CD69+CD4+ T cell (%)
ns
P=0.3054
Spearman r=0.1191
(n=76)
CD25+CD4+ T cell (%)
ns
P=0.1209
Spearman r=0.1794
(n=76)
PD-1+CD4+ T cell (%)
ns
P=0.0695
Spearman r=0.2094
(n=76)
TIM-3+CD4+ T cell (%)
ns
P=0.7502
Spearman r=0.0438
(n=77)
Gr.B+CD4+ T cell (%)
ns
P=0.6110
Spearman r=0.0589
(n=77)
CD69+CD8+ T cell (%)
ns
P=0.2711
Spearman r=0.1287
(n=75)
CD95+CD8+ T cell (%)
ns
P=0.8224
Spearman r=0.0262
(n=76)
CD25+CD8+ T cell (%)
ns
P=0.9685
Spearman r=-0.0046
(n=77)
CD40L+CD8+ T cell (%)
ns
P=0.0801
Spearman r=0.2020
(n=76)
PD-1+CD8+ T cell (%)
ns
P=0.2955
Spearman r=0.1216
(n=76)
CTLA-4+CD8+ T cell (%)
ns
P=0.3809
Spearman r=0.1012
(n=77)
CD107a+CD8+ T cell (%)
B
ns
P=0.1050
Spearman r=-0.1396
(n=136)
CD69+CD4+ T cell(%)
HBsAg($log_{10}$ IU/mL)
ns
P=0.3799
Spearman r=-0.0745
(n=141)
HLA-DR+CD4+ T cell(%)
ns
P=0.4892
Spearman r=-0.0591
(n=139)
CD25+CD4+ T cell(%)
ns
P=0.6412
Spearman r=-0.0400
(n=138)
CD95+CD4+ T cell(%)
ns
P=0.4673
Spearman r=-0.0617
(n=141)
CD40L+CD4+ T cell(%)
ns
P=0.9251
Spearman r=0.0081
(n=139)
PD-1+CD4+ T cell(%)
ns
P=0.7846
Spearman r=-0.0235
(n=138)
TIM-3+CD4+ T cell(%)
ns
P=0.5898
Spearman r=-0.0468
(n=135)
CTLA-4+CD4+ T cell(%)
ns
P=0.3007
Spearman r=-0.0878
(n=141)
Gr.B+CD4+T cells (%)
ns
P=0.9754
Spearman r=-0.0027
(n=138)
CD107a+CD4+ T cell(%)
ns
P=0.2823
Spearman r=-0.0929
(n=136)
CD69+CD8+ T cell(%)
ns
P=0.3664
Spearman r=-0.0767
(n=141)
HLA-DR+CD8+ T cell(%)
ns
P=0.5714
Spearman r=-0.0484
(n=139)
CD25+CD8+ T cell(%)
ns
P=0.8754
Spearman r=-0.0135
(n=138)
CD95+CD8+ T cell(%)
ns
P=0.0734
Spearman r=-0.1513
(n=141)
CD40L+CD8+ T cell(%)
ns
P=0.1139
Spearman r=0.1347
(n=139)
PD-1+CD8+ T cell(%)
ns
P=0.4655
Spearman r=0.0626
(n=138)
TIM-3+CD8+ T cell(%)
ns
P=0.7459
Spearman r=-0.0281
(n=135)
CTLA-4+CD8+ T cell(%)
ns
P=0.3344
Spearman r=-0.0819
(n=141)
Gr.B+CD8+T cells (%)
ns
P=0.4683
Spearman r=-0.0623
(n=138)
CD107a+CD8+ T cell(%)

C

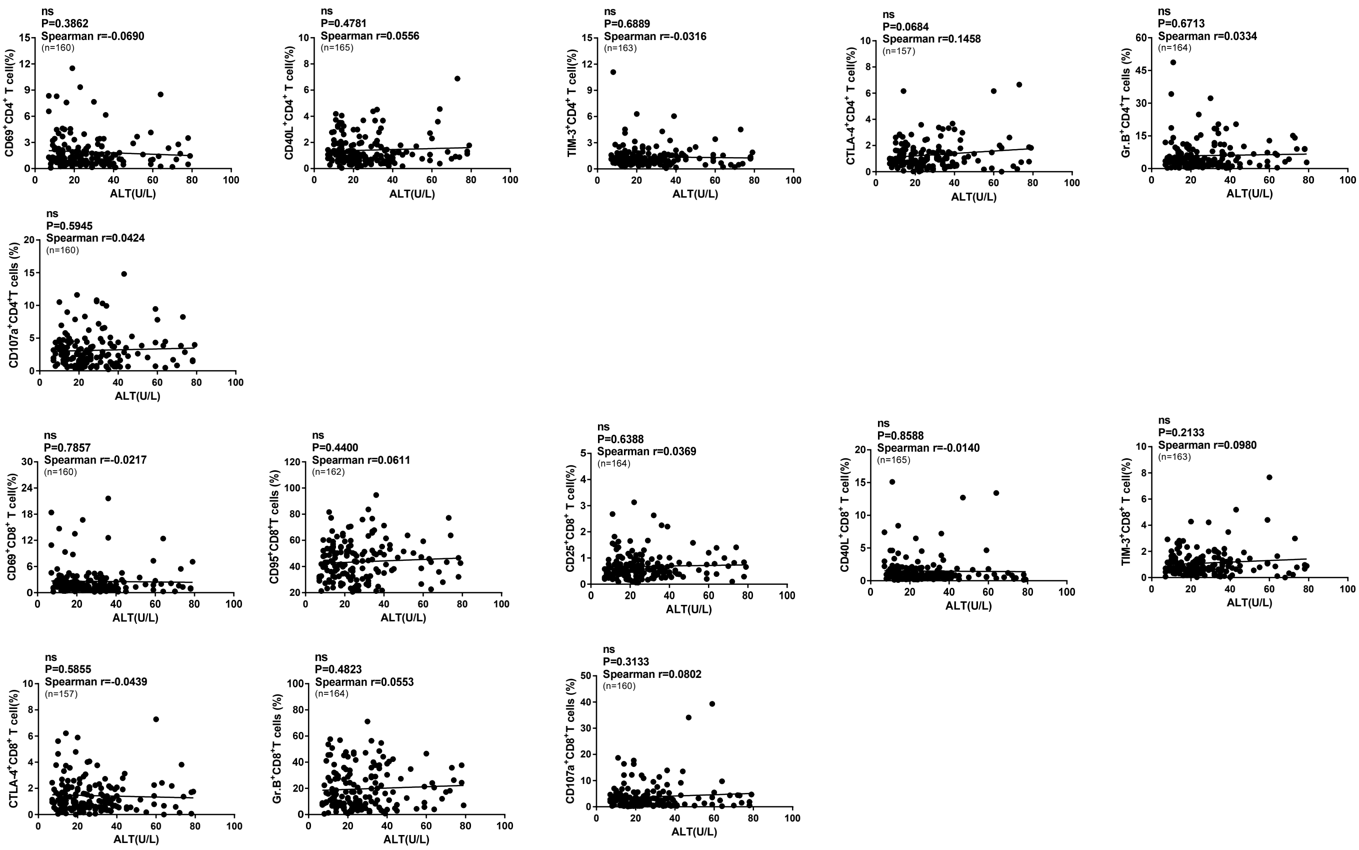
ns
P=0.3862
Spearman r=-0.0690
(n=160)
CD69+CD4+ T cell(%)
ALT(U/L)
ns
P=0.4781
Spearman r=0.0556
(n=165)
CD40L+CD4+ T cell(%)
ALT(U/L)
ns
P=0.6889
Spearman r=-0.0316
(n=163)
TIM-3+CD4+ T cell(%)
ALT(U/L)
ns
P=0.0684
Spearman r=0.1458
(n=157)
CTLA-4+CD4+ T cell(%)
ALT(U/L)
ns
P=0.6713
Spearman r=0.0334
(n=164)
Gr.B+CD4+T cells (%)
ALT(U/L)
ns
P=0.5945
Spearman r=0.0424
(n=160)
CD107a+CD4+T cells (%)
ALT(U/L)
ns
P=0.7857
Spearman r=-0.0217
(n=160)
CD69+CD8+ T cell(%)
ALT(U/L)
ns
P=0.4400
Spearman r=0.0611
(n=162)
CD95+CD8+T cells (%)
ALT(U/L)
ns
P=0.6388
Spearman r=0.0369
(n=164)
CD25+CD8+ T cell(%)
ALT(U/L)
ns
P=0.8588
Spearman r=-0.0140
(n=165)
CD40L+CD8+ T cell(%)
ALT(U/L)
ns
P=0.2133
Spearman r=0.0980
(n=163)
TIM-3+CD8+ T cell(%)
ALT(U/L)
ns
P=0.5855
Spearman r=-0.0439
(n=157)
CTLA-4+CD8+ T cell(%)
ALT(U/L)
ns
P=0.4823
Spearman r=0.0553
(n=164)
Gr.B+CD8+T cells (%)
ALT(U/L)
ns
P=0.3133
Spearman r=0.0802
(n=160)
CD107a+CD8+T cells (%)
ALT(U/L)

D

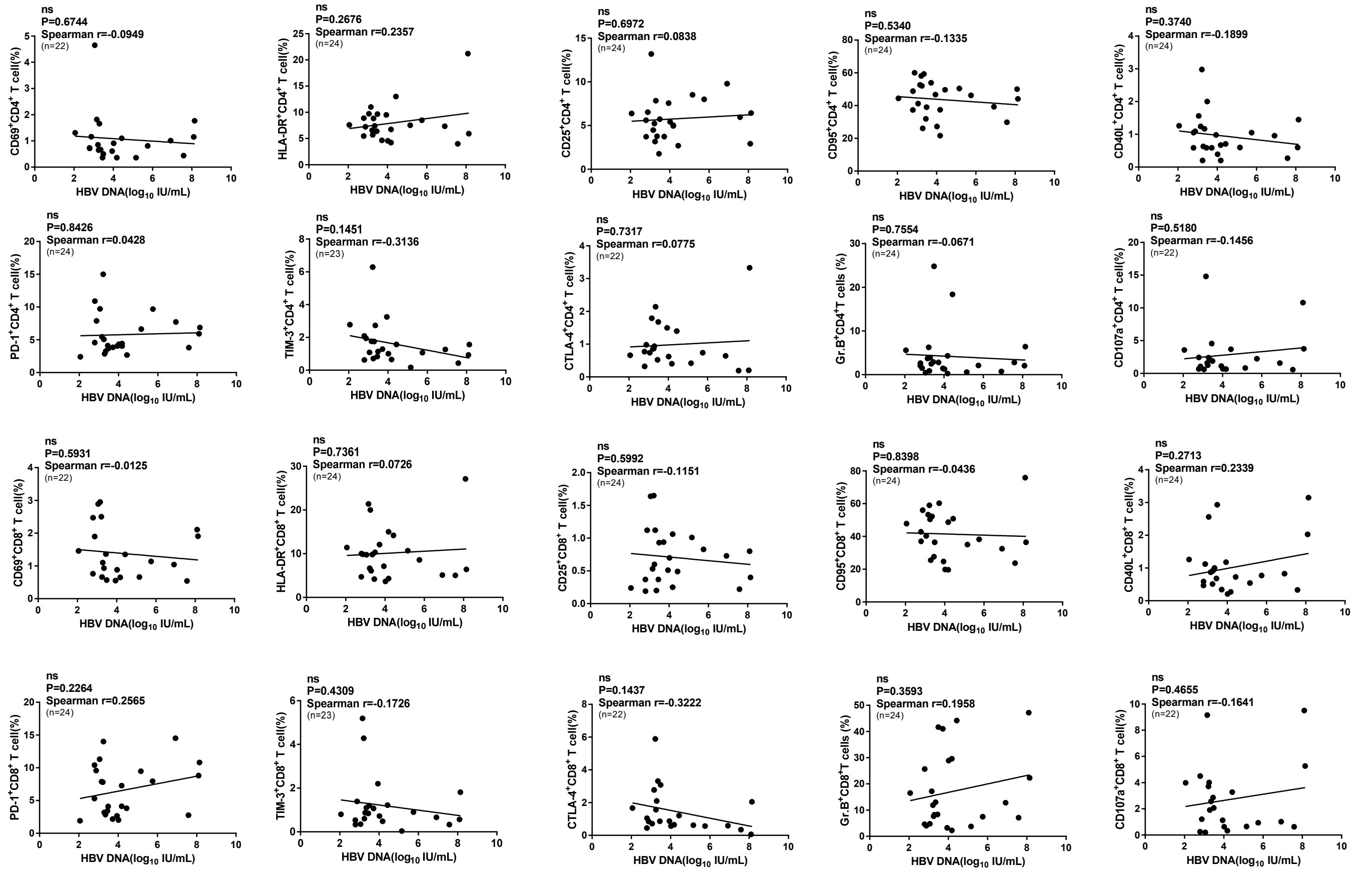
ns
P=0.6744
Spearman r=-0.0949
(n=22)
CD69+CD4+ T cell(%)
HBV DNA(log10 IU/mL)
ns
P=0.2676
Spearman r=0.2357
(n=24)
HLA-DR+CD4+ T cell(%)
HBV DNA(log10 IU/mL)
ns
P=0.6972
Spearman r=0.0838
(n=24)
CD25+CD4+ T cell(%)
HBV DNA(log10 IU/mL)
ns
P=0.5340
Spearman r=-0.1335
(n=24)
CD95+CD4+ T cell(%)
HBV DNA(log10 IU/mL)
ns
P=0.3740
Spearman r=-0.1899
(n=24)
CD40L+CD4+ T cell(%)
HBV DNA(log10 IU/mL)
ns
P=0.8426
Spearman r=0.0428
(n=24)
PD-1+CD4+ T cell(%)
HBV DNA(log10 IU/mL)
ns
P=0.1451
Spearman r=-0.3136
(n=23)
TIM-3+CD4+ T cell(%)
HBV DNA(log10 IU/mL)
ns
P=0.7317
Spearman r=0.0775
(n=22)
CTLA-4+CD4+ T cell(%)
HBV DNA(log10 IU/mL)
ns
P=0.7554
Spearman r=-0.0671
(n=24)
Gr.B+CD4+T cells (%)
HBV DNA(log10 IU/mL)
ns
P=0.5180
Spearman r=-0.1456
(n=22)
CD107a+CD4+ T cell(%)
HBV DNA(log10 IU/mL)
ns
P=0.5931
Spearman r=-0.0125
(n=22)
CD69+CD8+ T cell(%)
HBV DNA(log10 IU/mL)
ns
P=0.7361
Spearman r=0.0726
(n=24)
HLA-DR+CD8+ T cell(%)
HBV DNA(log10 IU/mL)
ns
P=0.5992
Spearman r=-0.1151
(n=24)
CD25+CD8+ T cell(%)
HBV DNA(log10 IU/mL)
ns
P=0.8398
Spearman r=-0.0436
(n=24)
CD95+CD8+ T cell(%)
HBV DNA(log10 IU/mL)
ns
P=0.2713
Spearman r=0.2339
(n=24)
CD40L+CD8+ T cell(%)
HBV DNA(log10 IU/mL)
ns
P=0.2264
Spearman r=0.2565
(n=24)
PD-1+CD8+ T cell(%)
HBV DNA(log10 IU/mL)
ns
P=0.4309
Spearman r=-0.1726
(n=23)
TIM-3+CD8+ T cell(%)
HBV DNA(log10 IU/mL)
ns
P=0.1437
Spearman r=-0.3222
(n=22)
CTLA-4+CD8+ T cell(%)
HBV DNA(log10 IU/mL)
ns
P=0.3593
Spearman r=0.1958
(n=24)
Gr.B+CD8+T cells (%)
HBV DNA(log10 IU/mL)
ns
P=0.4655
Spearman r=-0.1641
(n=22)
CD107a+CD8+ T cell(%)
HBV DNA(log10 IU/mL)

### Figure S6

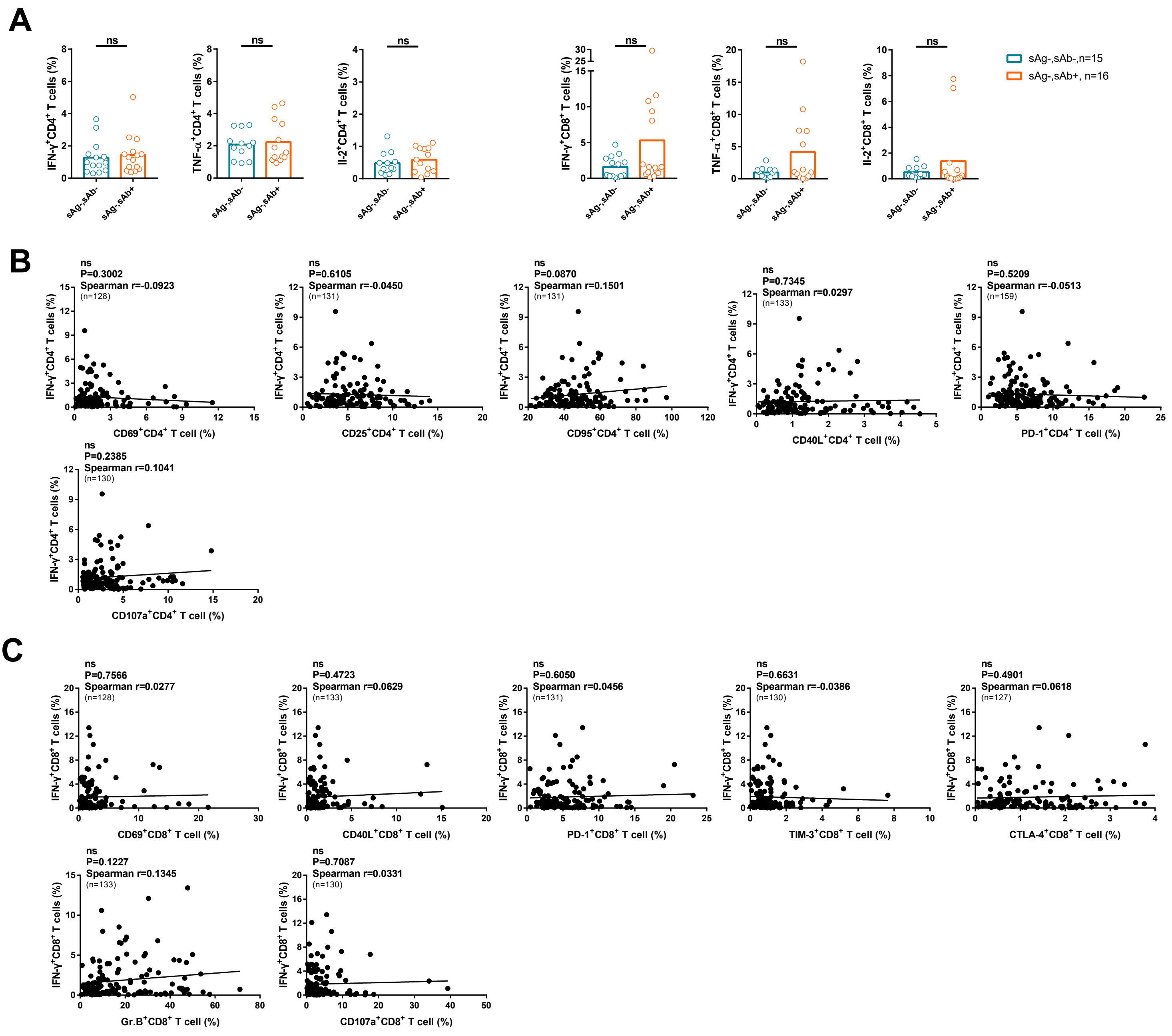
A
ns
IFN-γ+CD4+ T cells (%)
TNF-α+CD4+ T cells (%)
Il-2+CD4+ T cells (%)
IFN-γ+CD8+ T cells (%)
TNF-α+CD8+ T cells (%)
Il-2+CD8+ T cells (%)
sAg-,sAb-
sAg-,sAb+
sAg-,sAb-,n=15
sAg-,sAb+, n=16
B
ns
P=0.3002
Spearman r=-0.0923
(n=128)
CD69+CD4+ T cell (%)
ns
P=0.6105
Spearman r=-0.0450
(n=131)
CD25+CD4+ T cell (%)
ns
P=0.0870
Spearman r=0.1501
(n=131)
CD95+CD4+ T cell (%)
ns
P=0.7345
Spearman r=0.0297
(n=133)
CD40L+CD4+ T cell (%)
ns
P=0.5209
Spearman r=-0.0513
(n=159)
PD-1+CD4+ T cell (%)
ns
P=0.2385
Spearman r=0.1041
(n=130)
CD107a+CD4+ T cell (%)
C
ns
P=0.7566
Spearman r=0.0277
(n=128)
CD69+CD8+ T cell (%)
ns
P=0.4723
Spearman r=0.0629
(n=133)
CD40L+CD8+ T cell (%)
ns
P=0.6050
Spearman r=0.0456
(n=131)
PD-1+CD8+ T cell (%)
ns
P=0.6631
Spearman r=-0.0386
(n=130)
TIM-3+CD8+ T cell (%)
ns
P=0.4901
Spearman r=0.0618
(n=127)
CTLA-4+CD8+ T cell (%)
ns
P=0.1227
Spearman r=0.1345
(n=133)
Gr.B+CD8+ T cell (%)
ns
P=0.7087
Spearman r=0.0331
(n=130)
CD107a+CD8+ T cell (%)

### Figure S7

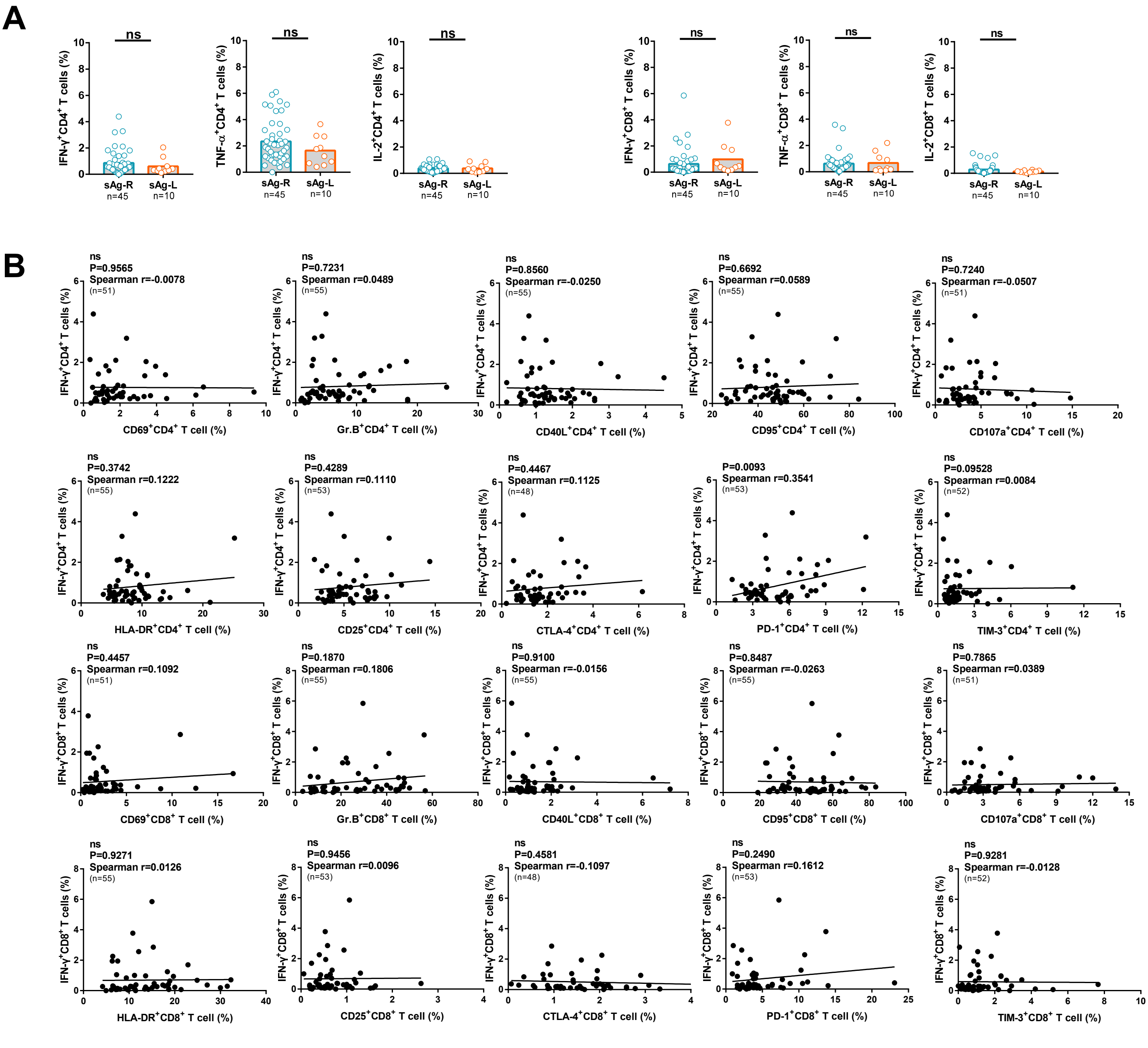
A
ns
IFN-γ+CD4+ T cells (%)
TNF-α+CD4+ T cells (%)
IL-2+CD4+ T cells (%)
IFN-γ+CD8+ T cells (%)
TNF-α+CD8+ T cells (%)
IL-2+CD8+ T cells (%)
sAg-R
n=45
sAg-L
n=10
B
ns
P=0.9565
Spearman r=-0.0078
(n=51)
CD69+CD4+ T cell (%)
ns
P=0.7231
Spearman r=0.0489
(n=55)
Gr.B+CD4+ T cell (%)
ns
P=0.8560
Spearman r=-0.0250
(n=55)
CD40L+CD4+ T cell (%)
ns
P=0.6692
Spearman r=0.0589
(n=55)
CD95+CD4+ T cell (%)
ns
P=0.7240
Spearman r=-0.0507
(n=51)
CD107a+CD4+ T cell (%)
ns
P=0.3742
Spearman r=0.1222
(n=55)
HLA-DR+CD4+ T cell (%)
ns
P=0.4289
Spearman r=0.1110
(n=53)
CD25+CD4+ T cell (%)
ns
P=0.4467
Spearman r=0.1125
(n=48)
CTLA-4+CD4+ T cell (%)
P=0.0093
Spearman r=0.3541
(n=53)
PD-1+CD4+ T cell (%)
ns
P=0.09528
Spearman r=0.0084
(n=52)
TIM-3+CD4+ T cell (%)
ns
P=0.4457
Spearman r=0.1092
(n=51)
CD69+CD8+ T cell (%)
ns
P=0.1870
Spearman r=0.1806
(n=55)
Gr.B+CD8+ T cell (%)
ns
P=0.9100
Spearman r=-0.0156
(n=55)
CD40L+CD8+ T cell (%)
ns
P=0.8487
Spearman r=-0.0263
(n=55)
CD95+CD8+ T cell (%)
ns
P=0.7865
Spearman r=0.0389
(n=51)
CD107a+CD8+ T cell (%)
ns
P=0.9271
Spearman r=0.0126
(n=55)
HLA-DR+CD8+ T cell (%)
ns
P=0.9456
Spearman r=0.0096
(n=53)
CD25+CD8+ T cell (%)
ns
P=0.4581
Spearman r=-0.1097
(n=48)
CTLA-4+CD8+ T cell (%)
ns
P=0.2490
Spearman r=0.1612
(n=53)
PD-1+CD8+ T cell (%)
ns
P=0.9281
Spearman r=-0.0128
(n=52)
TIM-3+CD8+ T cell (%)
IFN-γ+CD4+ T cells (%)
IFN-γ+CD8+ T cells (%)

### Figure S8

A

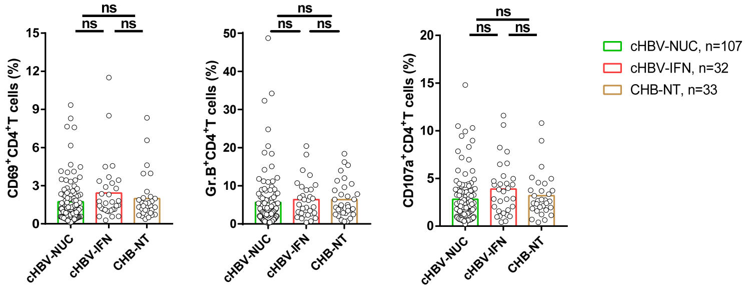

ns
CD69+CD4+T cells (%)
Gr.B+CD4+T cells (%)
CD107a+CD4+T cells (%)
cHBV-NUC
cHBV-IFN
CHB-NT
cHBV-NUC, n=107
cHBV-IFN, n=32
CHB-NT, n=33


B

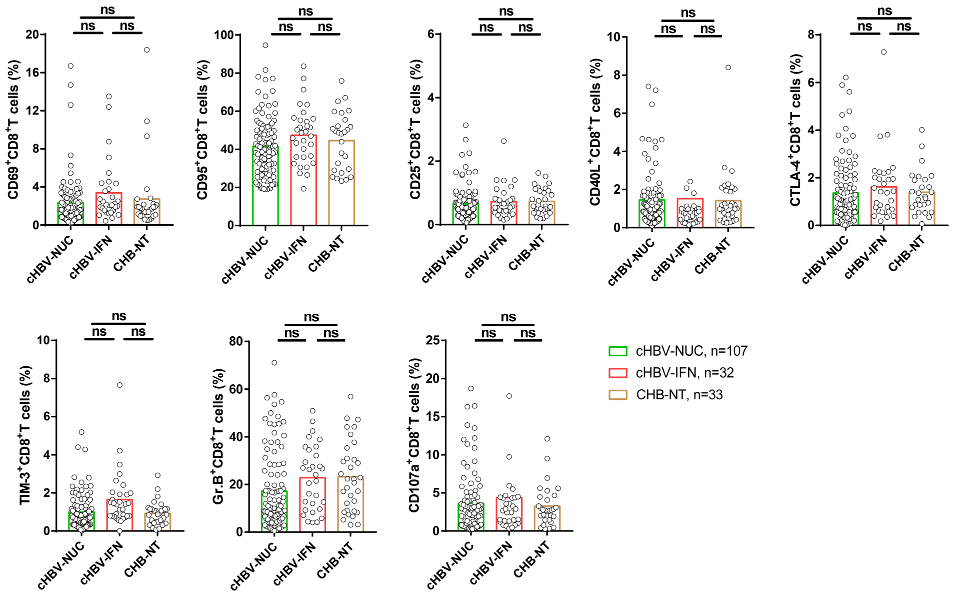

ns
CD69+CD8+T cells (%)
CD95+CD8+T cells (%)
CD25+CD8+T cells (%)
CD40L+CD8+T cells (%)
CTLA-4+CD8+T cells (%)
TIM-3+CD8+T cells (%)
Gr.B+CD8+T cells (%)
CD107a+CD8+T cells (%)
cHBV-NUC
cHBV-IFN
CHB-NT
cHBV-NUC, n=107
cHBV-IFN, n=32
CHB-NT, n=33

### Figure S9

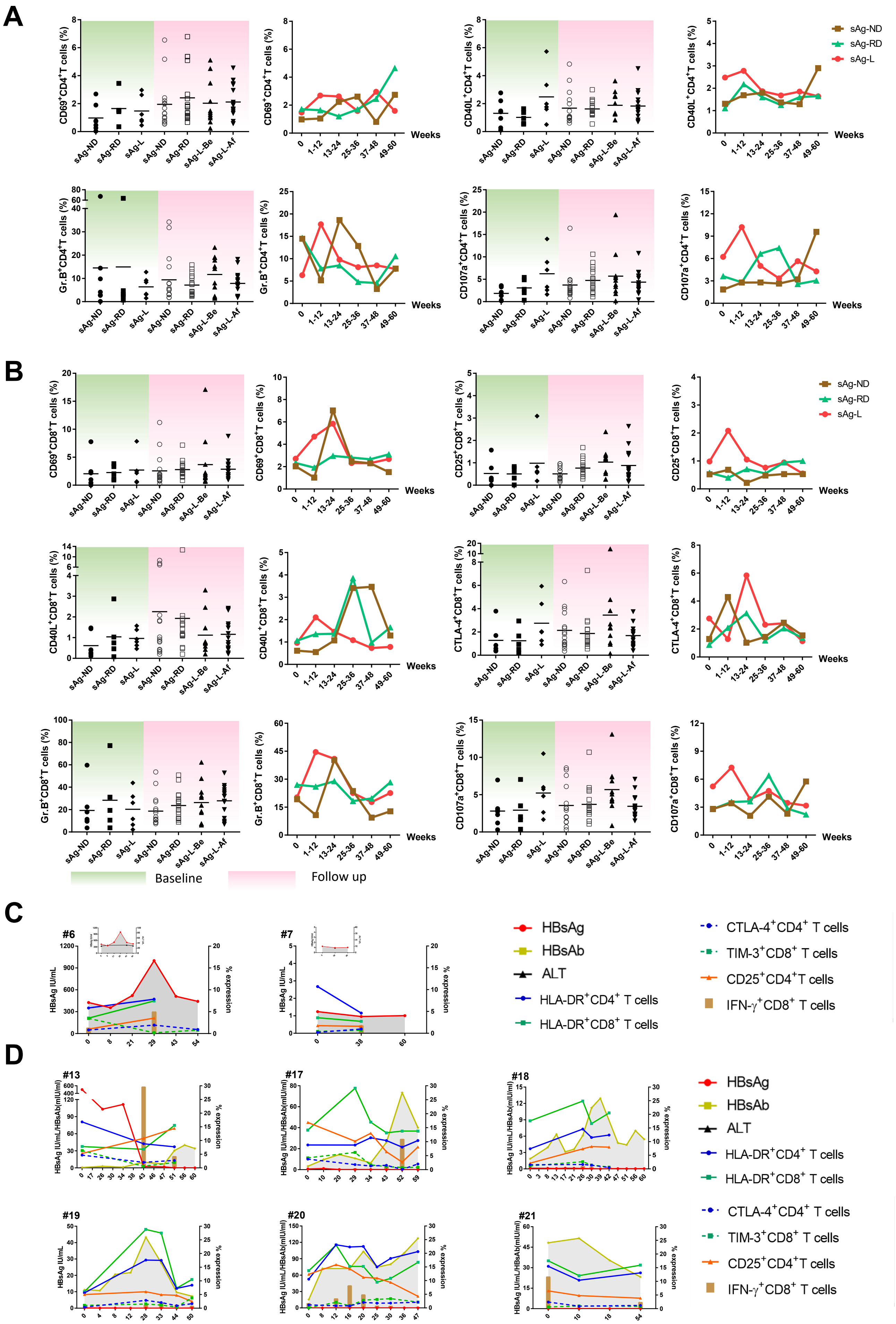

A
CD69+CD4+T cells (%)
CD40L+CD4+T cells (%)
Gr.B+CD4+T cells (%)
CD107a+CD4+T cells (%)
sAg-ND
sAg-RD
sAg-L
sAg-L-Be
sAg-L-Af
Weeks
0
1-12
13-24
25-36
37-48
49-60
B
CD69+CD8+T cells (%)
CD25+CD8+T cells (%)
CD40L+CD8+T cells (%)
CTLA-4+CD8+T cells (%)
Gr.B+CD8+T cells (%)
CD107a+CD8+T cells (%)
Baseline
Follow up
C
#6
#7
HBsAg IU/mL
% expression
HBsAg
HBsAb
ALT
HLA-DR+CD4+ T cells
HLA-DR+CD8+ T cells
CTLA-4+CD4+ T cells
TIM-3+CD8+ T cells
CD25+CD4+T cells
IFN-γ+CD8+ T cells
D
#13
#17
#18
#19
#20
#21
HBsAg IU/mL/HBsAb(mIU/ml)
HBsAg IU/mL
% expression
